# Rapid differentiation of regulatory CD4^+^ T cells in the infarcted myocardium blunts in situ inflammation

**DOI:** 10.1101/2022.03.25.485791

**Authors:** Murilo Delgobo, Emil Weiß, Diyaa ElDin Ashour, Lisa Popiolkowski, Panagiota Arampatzi, Verena Stangl, Paula Arias-Loza, Peter P. Rainer, Antoine-Emmanuel Saliba, Burkhard Ludewig, Ulrich Hofmann, Stefan Frantz, Gustavo Campos Ramos

## Abstract

**Background:** Myocardial infarction (MI) is a sterile inflammatory condition associated with tissue injury that results in the activation of T helper cell targeting cardiac antigens. However, the differentiation trajectories and *in situ* activity of heart-specific CD4^+^T cells activated in the MI context remain poorly understood.

**Methods:** Herein, we combined T-cell receptor transgenic models targeting myocardial protein, single-cell transcriptomics, and functional phenotyping to elucidate how the myosin-specific CD4^+^ T cells (TCR-M) differentiate in the murine infarcted myocardium and ultimately influence tissue repair. Furthermore, we adoptively transferred heart-specific T-cells that were pre-differentiated in vitro towards pro-inflammatory versus regulatory phenotypic states to dissect how they differentially regulate post-myocardial infarction (MI) inflammation.

**Results:** Flow cytometry and single-cell transcriptomics findings reveled that transferred TCR-M cells rapidly acquired an induced regulatory phenotype (iTreg) in the infarcted myocardium and blunt local inflammation. Myocardial TCR-M cells differentiated into two main lineages enriched with cell activation and pro-fibrotic transcripts (e.g. *Tgfb1*) or with suppressor immune checkpoints (e.g. *Pdcd1*), which we also found in human myocardial tissue. These cells produced high levels of latency-associated peptide (LAP) and inhibited interleukine-17 (IL-17) responses. Notably, TCR-M cells that were pre-differentiated in vitro towards a regulatory phenotype maintained a stable in vivo FOXP3 expression and anti-inflammatory activity when adoptively transferred prior to MI induction. In contrast, TCR-M cells that were pre-differentiated in vitro towards a pro-inflammatory T_H_17 phenotype were partially converted towards a regulatory phenotype in the injured myocardium and blunted myocardial inflammation.

**Conclusions:** These findings reveal that the myocardial milieu provides a suitable environment for iTreg differentiation and reveals novels mechanisms by which the healing myocardium shapes local immunological processes.

## Introduction

Research conducted over the past decade has positioned the immune system in the spotlight of cardiovascular biology, as immune phenomena mediate cardiac homeostatic functions and response to injury. Several immune cell types that contribute to electrical conduction, metabolism and tissue clearance in the healthy myocardium have been described ^1, 2^. During a stressful condition such as myocardial infarction (MI), necrotic cell death propels the rapid release of damage associated molecular patterns and autoantigens, resulting in inflammatory responses molded to bring back cardiac homeostasis^3^. Such responses are characterized by an early influx of neutrophils, followed by monocyte recruitment and then T- and B cell migration^1^. However, uncontrolled long-lasting immune responses can lead to adverse cardiac remodeling and further deteriorate cardiac function^1^. Understanding the cell types and kinetics involved in both scenarios is therefore critical for designing new therapies-and improving already existing ones and consequently promoting patient survival.

CD4^+^ T-cell-mediated responses have been shown to directly affect tissue repair in a wide range of animals and tissues, including the myocardium ^4-6^. Previous studies by our group and others demonstrated that CD4^+^ T-cells, particularly Tregs expressing the transcription factor FOXP3, can foster myocardial healing ^3, 5^. Increased T-cell signal was observed in heart-draining lymph nodes of infarcted patients and Tregs are present in cardiac biopsies ^7^, suggesting those responses are conserved in humans ^8^. Yet chronic T-cell activation has been shown to mediate detrimental remodeling during both pressure overload and aging ^2, 9^. In addition, bystander T-cell activation during viral infection and immune checkpoint inhibitor treatment can lead to myocarditis ^10, 11^. Defining CD4^+^ T-cell responses in the injured myocardium is thus crucial for proper therapeutic intervention.

Our previous study revealed that transferred T-cells specific to a myosin heavy alpha chain-derived peptide (MYHCA_614-629_), henceforth termed TCR-M cells, acquired FOXP3 expression when they reached the infarcted myocardium, a phenomenon which was associated with improved cardiac repair^7^. Tregs can exhibit broad phenotypic plasticity, though influenced by TCR signaling, tissue milieu and cell ontogeny, meaning they can influence the infarcted heart by varied and still poorly understood mechanisms^12^.

Tregs present in injured skeletal muscle and lung can promote tissue repair via a mechanism that depends on amphiregulin secretion; this significantly differs from the canonical suppression seen in experimental models of autoimmunity ^13, 14^. Under inflammatory conditions, Tregs may lose FOXP3 expression and acquire the pro-inflammatory phenotype characteristic of effector subsets. For instance, in the context of chronic ischemic heart failure, Bansal, et al. reported that Tregs acquire T_H_1 features and contribute to adverse left ventricular remodeling ^15^. Fate mapping experiments conducted in the context of atherosclerosis also revealed that Tregs specific to apolipoprotein B lose their suppressive capacity and shift towards a T_H_1/T_H_17 phenotype as atherogenesis progresses ^16^.

Considering the complex roles Tregs play in different disease models and stages ^17^, we performed deep phenotyping of TCR-M cells that engage post-MI responses in order to functionally dissect how different T-helper cell states regulate myocardial repair. By combining adoptive cell transfer models using cardiac antigen-specific transgenic CD4^+^ T-cells with single-cell transcriptomic and functional characterization, our analysis revealed a rapid differentiation of myosin-specific regulatory CD4^+^ T cells in the infarcted myocardium, which then blunted in situ inflammation and preserved cardiac functionality. The myocardial induced Tregs exhibited two main transcriptional states, one enriched with effector/ pro-fibrotic transcripts and the other enriched with suppressor immune checkpoints, which we also found in human myocardial tissue. Moreover, adoptive cell transfer of TCR-M cells previously polarized towards regulatory vs. pro-inflammatory states revealed the differential contribution of each major T-cell subset to the regulation of myocardial inflammation.

## Methods

### Data availability

The full methods and supplemental figures are available in the Supplemental Material. The raw transcriptomic data acquired in this study will be available after the peered reviewed version is published.

### Mice

Thy1.2 BALB/c mice were purchased form Charles River (Sulzfeld, Germany) and housed under specific pathogen-free conditions throughout the experiments. Thy1.1TCR-M mice expressing a transgenic TCR against MYHCA_614-629_ peptide presented on I-A^d^ (C.CB6-Tg(*Tcra,Tcrb)*^*562Biat*^ were bred in our housing facility^18^ and used as donors in adoptive T-cell transfer experiments. Thy1.2 DO11.10 mice expressing a transgenic TCR against Ovalbumin_323-339_ were housed under specific pathogen-free conditions and bred in our housing facility. All mouse strains share the same genetic background (BALB/c).

### Experimental models

Magnetically sorted Thy1.1 TCR-M CD4^+^ T-cells were adoptively transferred into Thy1.2 WT BALB/c and in DO11.10 mice (5 ×10^6^ cells > 90% purity, intraperitoneally injected) one day prior to induction of experimental myocardial infarction via permanent ligation of the left coronary artery. TCR-M cells found in the dissociated heart tissue, spleens and heart-draining mediastinal lymph nodes of infarcted recipients were immunophenotyped on days 5 and 7 after surgery by flow cytometry and single-cell RNA sequencing (see below). Echocardiography imaging of mice under slight isoflurane anesthesia (0.5-1.5%) was performed on day 5, as previously described ^19^, to assess infarct sizes and cardiac function following cell transfer experiments. The echocardiography imaging analyses was conducted by an experimenter blinded to cohort details. Animals with infarct size smaller than 20% were considered technical failures and excluded from analyses ^20^.

### Single-cell RNA sequencing

Collagenase-digested hearts and mechanically dissociated mediastinal lymph nodes were processed and stained with different combinations of hashtag TotalSeqC antibodies and CD90.1 CiteSeq antibody (clone OX-7). Live lineage negative (CD11b, CD8a, B220) CD4^+^ T-cells were sorted and combined as a single sample for library preparation using the 10x platform. Analysis was conducted using Seurat, scRepertoire and Monocle 3 R packages. A detailed description is found in Supplemental Materials.

### Statistical analyses

The results are shown as the mean ± the standard error of the mean (SEM) along with the distribution of all individual values in each group. Sample sizes for each group are described in figure legends. Graphs and statistical analyses were performed with GraphPad Prism (version 7.0, GraphPad Software, San Diego, CA, USA). Unpaired two-tailed *t-*test was used to compare two groups with data following normal distribution. For multiple comparisons between more than two groups, one or two-way analyses of variance (ANOVA) were conducted followed by *post hoc* test. Differences were considered significant for P values below 0.05.

### Study approval

The local authorities (Regierung von Unterfranken, Würzburg, Germany) approved all animal procedures and experiments were performed according to the Federation for Laboratory Animal Science Associations (FELASA) guidelines ^21^. Left ventricular myocardial tissue was obtained from patients deceased after myocardial infarction that underwent autopsy. The use of human tissue is conformed with legal and institutional requirements and was approved by the ethics committee of the Medical University of Graz (31-288 ex 18/19).

## Results

### Transferred heart-specific CD4^+^ T-cells acquire a regulatory phenotype in the heart and mediastinal lymph nodes

MI leads to cardiomyocyte death and subsequent release of autoantigens that stimulate T-cell responses. We have previously identified a peptide fragment derived from the cardiac myosin heavy alpha chain (MYHCA_614-629_) as the dominant antigen triggering post-MI CD4^+^ T-cell responses in BALB/c mice^7^. Adoptively transferred T-cells expressing a transgenic T-cell receptor specific for this myosin antigen (TCR-M) accumulated in the heart and acquired FOXP3 expression (**Figure 1A**). To determine whether local TCR activation is required for regulatory polarization in the heart, OVA-specific CD4^+^ T-cells obtained from DO11.10 mice were either pre-stimulated *in vitro* with their cognate antigen (OVA_323-339_) or kept in resting condition before being transferred into DO11.10 recipients one day prior to MI induction. Pre-activated but not resting DO11.10 cells accumulated in the infarcted heart and over 60% of the heart-infiltrating T helper cells expressed FOXP3, suggesting Treg conversion (**Figure 1B**). These findings indicate that TCR-dependent stimulation is required for T helper cells to infiltrate the injured heart ^22^ but dispensable for *in situ* Treg conversion.

**Figure 1.**
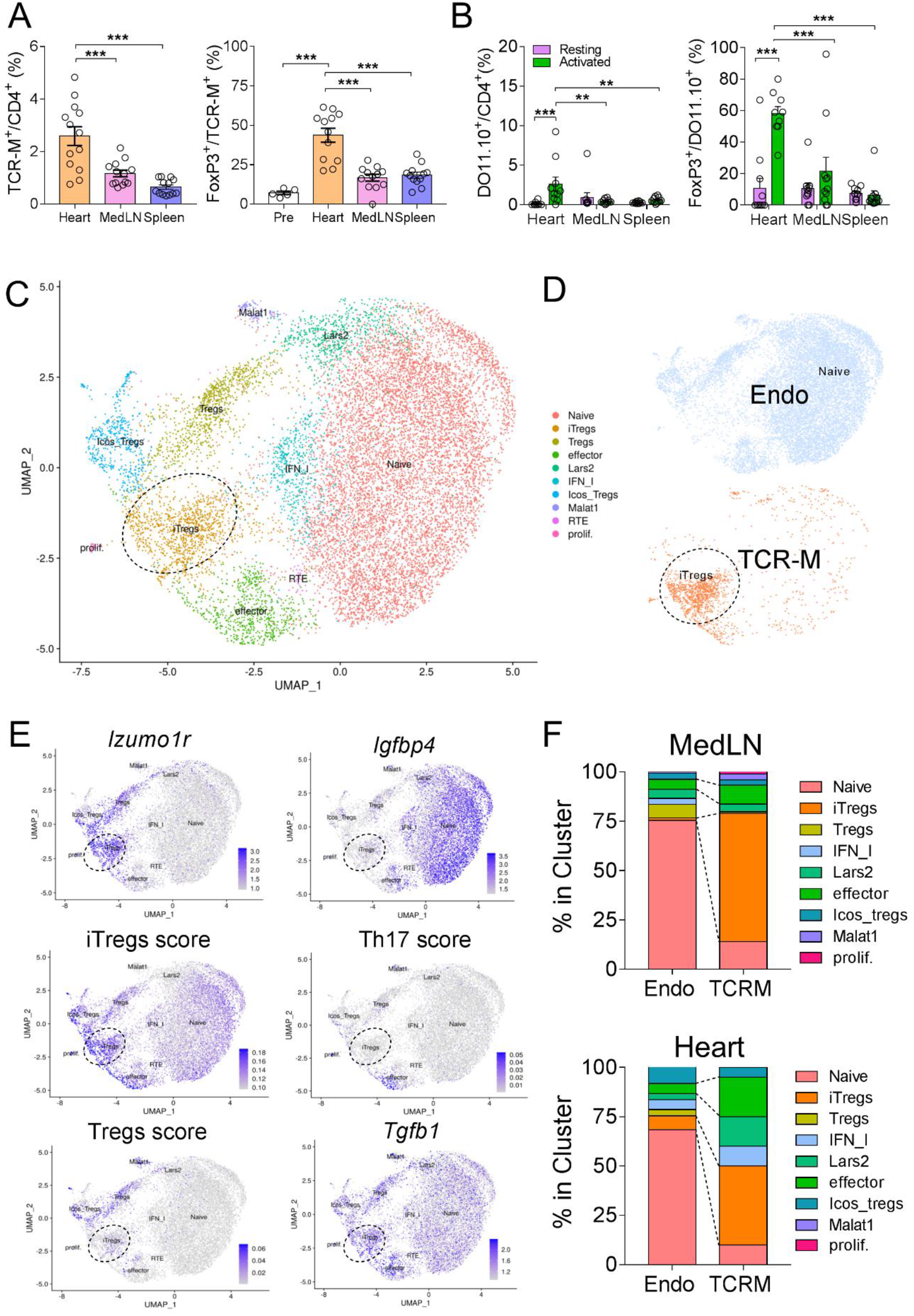
TCR-M cells shift towards an induced regulatory phenotype in the heart and mediastinal lymph nodes. **(A)** TCR-M cell frequency among endogenous CD4^+^ T-cells in heart, MedLN and spleen 7d after MI (left panel). Frequency of cardiac, MedLN and spleen TCR-M FOXP3^+^ cells at 7d after MI (right panel). **(B)** Frequency of resting (violet) and pre-activated (green) DO11.10 cells 5d after MI in DO11.10 recipients at different sites. Resting and pre-activated DO11.10 FOXP3^+^ cell distribution at different sites 5d after MI. **(C)** scRNA-seq analysis of total CD4^+^ T-cells from heart and MedLN of sham-operated and infarcted mice, 5 and 7d after surgery. UMAP representation of CD4^+^ T-cells based on *k*-nearest neighbor (KNN) cell clusters, identified by prototypic transcript expression. **(D)** UMAP representation of endogenous (blue) and TCR-M (orange) cells according to CD90.1.TotalSeqC expression (positive in TCR-M cells). **(E)** *Featureplots* depict the combined expression of cluster-defying markers and prototypic T_H_ gene sets in endogenous and TCR-M cells from Figure 1C. Dashed-line circles highlight the TCR-M cluster. **(F)** Cell numbers per scRNAseq clustering of endogenous and TCR-M cells in medLN (top) and heart (bottom) 5d after MI. Bars are color-coded according to Figure 1C and dashed lines indicate *iTreg* and effector cluster shifts for endogenous and TCR-M cells at each site. Panels **A-B** display the group mean values (bars); the error represents SEM and the circles the distribution of each individual value. Data were acquired from at least two independent experiments, n=5-12 mice. Statistical analysis in **A**: One-way ANOVA followed by Tukey’s post-test. *P < 0.05, **P < 0.01 and ***P<0.001. Statistical analysis in panel **B**: 2-way ANOVA followed by Sidak’s multiple comparisons test.

To assess in greater depth how the MI milieu favors regulatory phenotype development in cardiac-specific CD4^+^ T-cells, we performed single-cell RNA and T-cell receptor sequencing of endogenous CD4^+^ T-cells (singlets, live, CD45^+^, Lin^-^, CD4^+^, Thy1.1^-^) and TCR-M cells (singlets, live, CD45^+^, Lin^-^, CD4^+^ Thy1.1^+^) purified from hearts and mediastinal lymph nodes at 5 and 7 days after MI (**Suppl. Figure 1A**). All sorted cell subsets and groups were multiplexed using barcoded hashtag antibodies (anti-CD45/MHC-I TotalSeqC) and sequenced as a single library preparation to avoid batch effects (**Suppl. Figure 1A**). In total, 13,940 CD4^+^ single cells were analyzed and TCR-M cells were identified based on Thy1.1 Cite-seq signal in addition to TCR chain analysis, resulting in 1,565 TCR-M single cells (**Suppl. Figure 1A and 1B**). As illustrated in **Figure 1C**, our analyses revealed ten distinct clusters of cardiac and mediastinal lymph node T-cells, including *naïve* cells (expressing *Igfbp4, Il7r* and *Ccr7*), *bona fide Tregs* (expressing *Foxp3, Il2ra* and *Ctla4*) and effector T-cells (expressing *Tnfrsf9, Nr4a1* and *Cd69*^*variable*^), amongst others (**Figure 1C and Suppl. Figure 2A and 2B**). In addition, we identified a T-cell cluster, composed mainly of TCR-M cells, showing high expression of checkpoint inhibitor receptors (*Pdcd1, Cd200, Lag3*), regulatory markers (*Il2rb, Tgfb1*) and TCR activation genes (*Cd5, Nfatc1*) (**Figure 1C-1E and Suppl. Figure 2A, 2B**). We also identified five clusters of cells expressing type I interferon inducible genes, named *IFN_I* (*Ifit1, Isg15, Irf7* and *Stat1*); *Lars2* and *Malat1* clusters expressing mitochondrial genes and some TNF-responsive genes, respectively; two minor clusters of cycling T-cells (*prolif*.) expressing cyclin genes and a set of transcripts suggesting recent thymic emigrants *(RTE*) (**Figure 1C, Suppl. Figure 2A**).

TCR-M cells showed high expression levels of *Izumo1r* but lacked *Igfbp4* (**Figure 1E**), consistent with a previously established Treg signature ^23^. To further dissect the phenotype of TCR-M cells, we compiled a gene-set module score based on a RNA sequencing atlas ^24^ (**extended table 1**) of distinct polarized T-cell populations. As shown in **Figure 1E**, the TCR-M cells exhibited elevated expression of a transcriptomic signature compatible with “induced Tregs” (*iTregs*) phenotype ^24^, such as *Cd200, Pou2f2* and *Sox4*. TCR-M cells showed negligible expression of transcripts related to T_H_17 subset signature (**Figure 1E**). Despite the clear induced Treg signature, no *Foxp3* expression among TCR-M cells was detected at the mRNA level (**Suppl. Figure 2B**). This finding is compatible with the low FOXP3 expression level observed at the protein level (**Suppl. Figure 2C**) and the low sequencing depth inherent to this technology. Integrating our results with another available dataset on polyclonal cardiac T-cells (without known antigen specificity) ^25^ further confirmed that the TCR-M cells clustered together with an independent cardiac Treg cluster, providing evidence for their regulatory phenotype (**Supplemental Figure 3A**).

Analyses of cell cluster distributions indicated that over 65% of TCR-M cells in the MedLN exhibited an *iTreg* signature, while the second most represented clusters were *naïve* (14%) and *effector* (10%) cells (**Figure 1F**). Cardiac TCR-Ms showed more heterogeneous distribution, with 40% clustering as *iTregs* and 20% as *effector* cells (**Figure 1F**). Taken together, the flow cytometry and scRNAseq analyses shown in **Figure 1A-1F** suggested that myosin-specific CD4^+^ T-cells acquire an induced regulatory phenotype in the heart and MedLNs, a phenotype, which overlaps with endogenous *bona fide* Tregs, conventional effector cells and small pro-inflammatory cells.

### TCR-M cells activated in the context of MI present effector and suppressor phenotype signatures

To understand TCR-M cells’ transcriptomic signature in greater depth, we subset and re-clustered them for further analyses. We identified three main clusters in which TCR-M cells split, namely “*early/T*_*CM*_”, “*effector Tregs*” and “*suppressor*” (**Figure 2A**). The first cluster was comprised of cells exhibiting a mixed signature of early-activation transcripts and central memory phenotype, which largely overlap due to the similar circulation patterns exhibited by these two subsets (e.g. *Ccr7* and *Sell*) ^26^. The cluster termed “*effector Tregs*” encompassed cells marked by the expression of canonical T-cell activation genes (*Nr4a1, Tnfrsf9*), the transcription factor *Myc* combined with high expression levels of *Tgfb1* and an induced Treg signature (**Figure 2B**). The “*suppressor*” cluster was defined by the expression of genes encoding immune inhibitory receptors, especially *Pdcd1* encoding PD-1, **Figure 2C and Suppl. Figure 4A and 4B**). Myocardial left ventricular tissue of a patient that had died upon myocardial infarction confirmed the accumulation of lymphocytes (CD3^+^) in the scar tissue, alongside with cells expressing PD-1^+^ and TGF-β^+,^ the two main transcripts observed in our sequencing analyses (**Figure 2D**).

**Figure 2.**
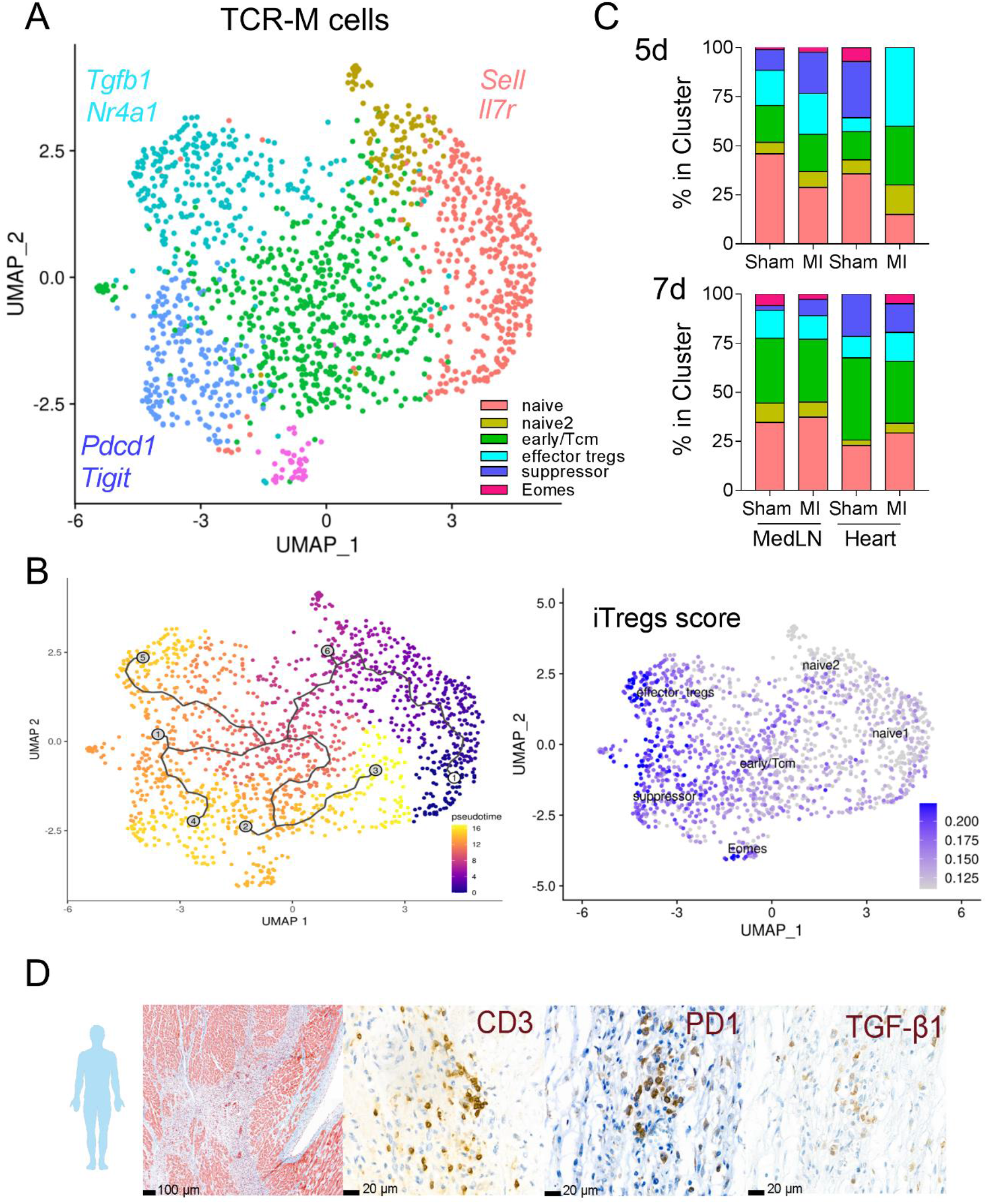
Phenotypic landscape of TCR-M cells in heart and MedLN of infarcted mice. **(A)** UMAP representation and re-clustering restricted to TCR-M cells from Figure **1D**. Colors depict newly classified clusters based on prototypic gene signatures and most expressed transcripts. **(B)** *Pseudotime* analysis (left panel) shows paths and degrees of differentiation. Cluster *naive1* was set as starting condition. Numbered nodes represent differentiation states and scale shows degree of differentiation. *Featureplot* illustrates combined *iTregs* signature scores for different TCR-M clusters (bottom). **(C)** Cell numbers per scRNAseq clustering of distinct TCR-M cell clusters from the MedLN and heart 5d (top) and 7d (bottom). Bars are color coded according to Figure 2A. **(D)** Analyses of a post-MI human heart, including Masson’s trichrome staining and immunohistochemistry demonstrate interstitial fibrosis (blue), inflammation and co-localization of T-cells (CD3), PD-1 and TGF-β1 in these areas.

To understand how TCR-M cells differentiate towards each state, we performed a *pseudotime* analysis, setting the *naïve1* cluster as an undifferentiated starting point (**Figure 2B, left panel**). Our analysis revealed that TCR-M cells reached an early activation state and then mainly branched into either *effector Tregs* (node 5) or *suppressor* (node 4) states (**Figure 2B**). Notably, as the TCR-M cells differentiated, they built up an *iTreg* signature (**Figure 2B, right panel, Suppl. Figure 4C**) which was retained in all activated states.

Next, we assessed the distribution of each TCR-M cluster in the MedLNs and hearts of sham-operated or infarcted mice. At day 5, TCR-Ms showing a *suppressor* phenotypic state were more frequent in the MedLNs of infarcted mice (20%) compared to sham controls (10%). Further, TCR-Ms found in infarcted hearts on day 5 mainly presented an *effector Treg* phenotype (40%) that was absent from sham-operated hearts, which mainly harbored TCR-M cells with a naïve phenotype (**Figure 2C**). On day 7 post MI, though, the TCR-M cells exhibited similar phenotypic distributions in the MedLNs and hearts of sham and infarcted mice (**Figure 2C**). Taken together, these findings show that the infarcted myocardium positions TCRM-cells in the injury site to acquire an *effector Treg* signature on day 5 post-injury. Moreover, that Treg conversion was also observed in sham-operated mice on day 7 post MI provides functional evidence that mechanisms of peripheral tolerance to cardiac antigens continuously operate during steady-state conditions, even in the absence of myocardial damage. These findings are particularly relevant for an autoantigen that is not expressed in the thymus and hence is not covered by central tolerance mechanisms ^27^.

To further explore how MI shapes TCR-M cell differentiation, we analyzed the expression of key transcripts and effector molecules in T-cell biology. As shown in **Supplementary figure 5A**, MedLN TCR-M cells purified from infarcted mice showed robust upregulation of genes associated with Treg fitness/suppressive function ^28, 29^ (e.g. *Ikzf2, Batf*), effector/memory responses (e.g. *Cd44, Tnfrsf9*) ^30^ and the cytokines *Tgfb1* and *Tnfsf8* at day 5 post MI (**Suppl. Figure 5A**). Myocardial TCR-M responses showed pronounced upregulation of effector molecules, the α4β1 integrin pair (*Itga4, Itgb1*) and downregulation of checkpoint inhibitors (*Cd96, Lag3*) (**Suppl. Figure 5B**). Day 7 MedLN responses were still characterized by a regulatory profile, but with downregulated effector molecules and higher expression of the cytokine *Il16*, which has been shown to mediate preferential Treg migration ^31^ (**Suppl. Figure 5A**). Similarly, day 7 cardiac responses showed increased levels of regulatory and pro-healing genes (*Il10ra, Tgfb1*) and upregulation of checkpoint inhibitors (**Suppl. Figure 5B**). Altogether, our data indicate that, in infarcted recipients, TCR-M cells differentiate towards an effector regulatory T-cell phenotype, peaking on day 5 post MI, and then shift towards a suppressive/pro-resolution profile at day 7.

### TCR-M cells foster a pro-healing program in fibroblasts/macrophages

We next sought to investigate how TCR-M-mediated responses occur in the heart and MedLNs, focusing on connections between cytokines, chemokines and receptors expressed by distinct TCR-M subsets and the respective receptors expressed by fibroblasts, macrophages and endothelial cells. To explore this, we integrated the transcripts up-regulated after MI in each of the TCR-M subsets defined in **Figure 2A** with a single-cell atlas consisting of several cell subsets purified from infarcted hearts 3 and 7 days post injury ^32^. TCR-M cells exhibiting *early/T*_*CM*_, *effector Tregs* and *suppressor* phenotypes were taken as seeder cells, while fibroblasts, macrophages and endothelial cells were taken as receiver cells. Only transcripts up-regulated by the MI condition were considered in this approach. *Nichenet* analysis revealed synergistic pathways induced by *suppressor* and *effector Tregs* TCR-M cells. Transcripts enriched in *effector Tregs* TCR-Ms showed crosstalk to molecules in fibroblasts related to tissue repair processes, such as “response to wounding” and “collagen fibril organization” (**Figure 3A-B**). In macrophages, *suppressor* cells promoted pro-survival (*Pim1)* and immunoregulatory genes (*Arg1, Il1rn* and *Ptgs2*) via *Il4, Tnfsf11* and *Il21* signaling. In addition, *effector Tregs* stimulated pro-survival, inflammatory responses and a pro-healing phenotype via *Tgfb1* and, to a lesser extent, *Tnf* (**Figure 3C-D**). Analyzing TCR-M-derived ligands against receptors expressed on endothelial cells after MI revealed biological processes related to stress response and mesenchymal transition (**Figure 3E-F**).

**Figure 3.**
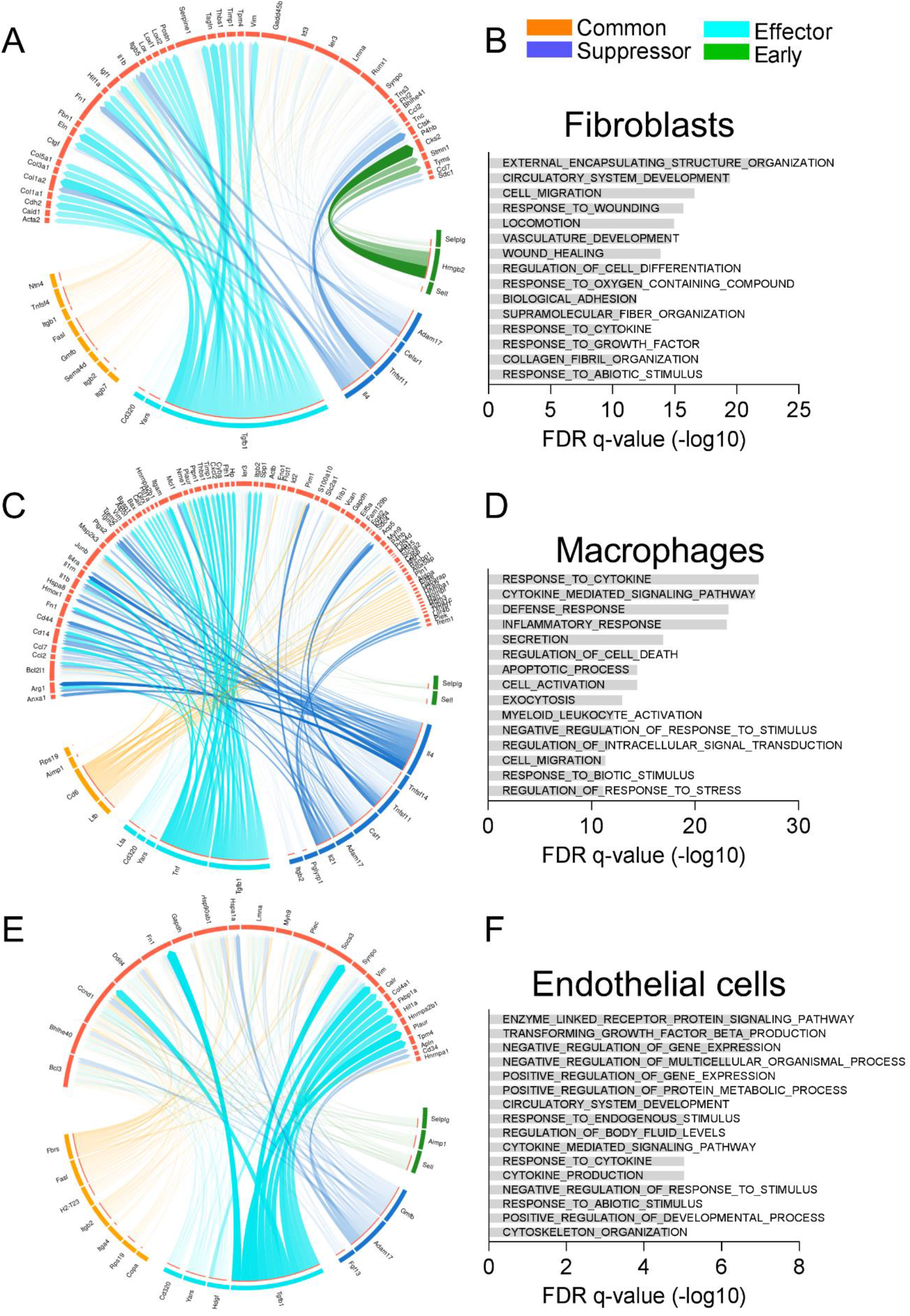
Nichenet analysis revealed synergistic pathways induced by TCR-M cells. **(A)** Circos plots illustrate fibroblast targets induced by TCR-M ligands per cell cluster. **(B)** Gene ontology analysis of transcripts induced in fibroblasts by TCR-M cells. **(C)** Circos plots show macrophage targets induced by TCR-M ligands and **(D)** gene ontology analysis of corresponding molecules. **(E)** Circos plots show TCR-M ligands and molecules induced in endothelial cells according to cell cluster (suppressor, effector, early or common mediator). **(F)** Gene ontology analysis of induced transcripts in endothelial cells. Panels **B, D** and **F** represent the top 15 gene ontology processes induced by TCR-M cells. The X-axes show the negative log10 FDR value.

### TCR-M cells activated in the context of MI suppress pro-inflammatory T_H_17 responses

To scrutinize the phenotype of myocardial T-cells at a functional level, we assessed the intracellular expression of prototypical T_H_ cytokines produced by TCR-M and endogenous CD4^+^ T-cells purified from infarcted hearts and spleen and re-stimulated in vitro (**Figure 4A**). We found that cardiac TCR-M cells preferentially expressed the latency associated peptide (LAP, a readout for TGF-β1 production), while showing low expression levels of pro-inflammatory mediators such as IFN-γ, IL-17 and TNF in accordance to the scRNAseq analysis (**Figure 4B**). In WT infarcted mice that have not received adoptive transfer of TCR-M cells, the endogenous cardiac CD4^+^ T-cell compartment was marked by a mixed expression of LAP and IL-17. However, the adoptive transfer of TCR-M cells suppressed the IL-17 production by the recipients’ endogenous CD4^+^ T-cells (**Figure 4B**). These findings provide a functional confirmation that TCR-M cells can suppress endogenous T-cell responses and T_H_17 polarization in the infarcted heart, as suggested by the transcriptomic signatures. Importantly, TCR-M cells found in the spleens of infarcted mice neither show a LAP-expressing signature nor suppressed the activity of neighboring endogenous T-cells (**Figure 4C**). Interestingly, the endogenous cardiac CD4^+^ T-cells of TCR-M transferred mice also showed a higher frequency of FOXP3^+^-expressing Tregs, in contrast to control mice (**Figure 4D**). No differences in the endogenous Treg numbers were found in the spleen, though (**Figure 4E**). These results illustrate that transferred TCR-M cells orchestrate local immune response by favoring Treg expansion/migration and inhibiting T_H_17 responses.

**Figure 4.**
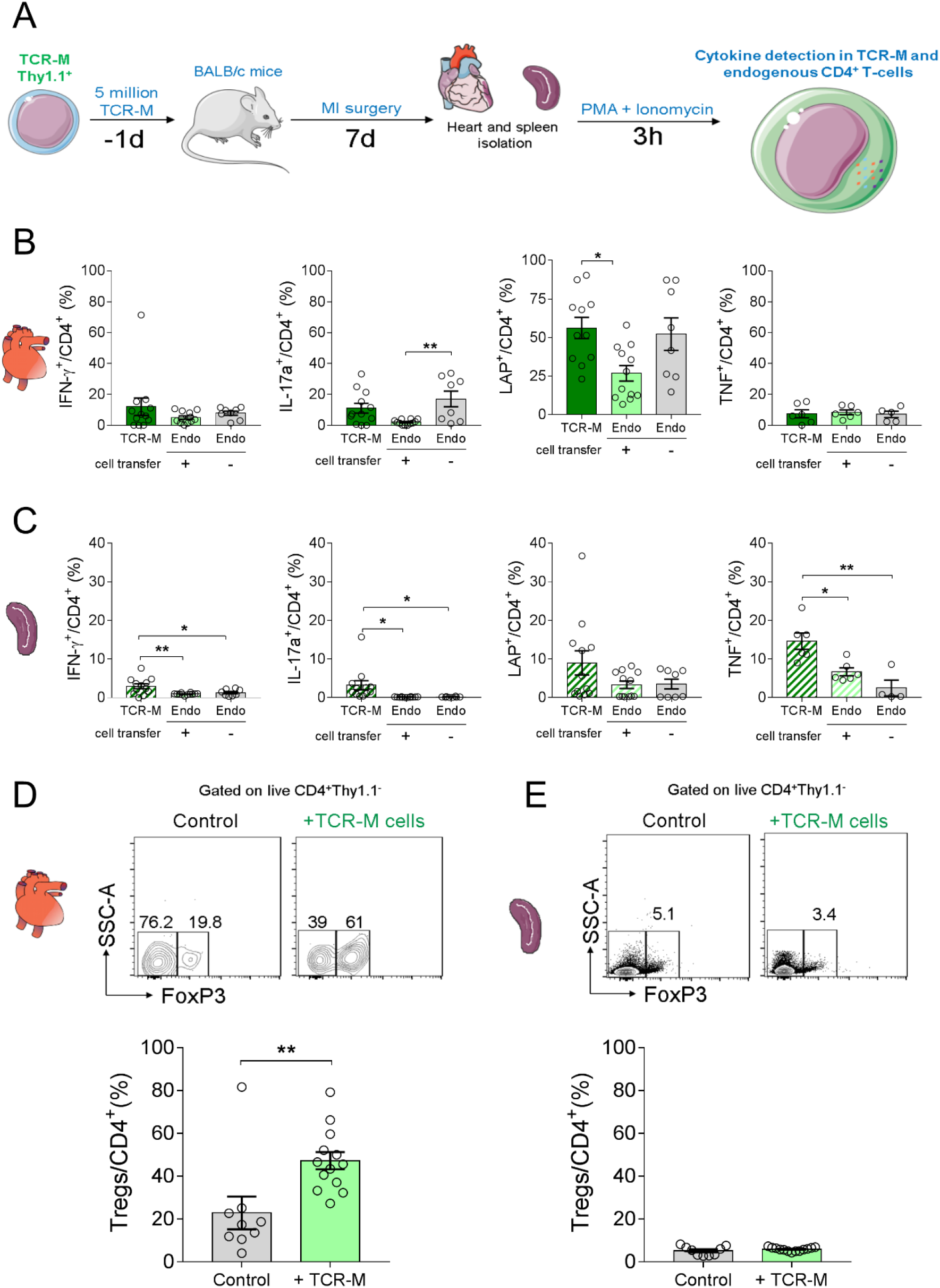
Cardiac TCR-M cells produce LAP and suppress IL-17 responses in endogenous CD4^+^ T-cells. **(A)** Experiment design: TCR-M cells were transferred to WT BALB/c mice one day prior to MI surgery. Hearts and spleens were collect 7d post MI and cells were stimulated for 3h with PMA/Ionomycin. Cytokine production was analyzed in TCR-M (CD4^+^ CD90.1^+^) and endogenous (CD4^+^CD90.1^-^) cells from TCR-M-transferred and control no-transfer mice. **(B)** Frequency of cardiac CD4^+^ IFN-γ, IL-17, LAP and TNF-producing cells in TCR-M cells (dark green), endogenous cells from TCR-M-transferred mice (light green) and endogenous cells from control no-transfer (grey) mice. **(C)** Corresponding intracellular cytokine analysis of spleens from TCR-M transferred and control no-transfer mice. **(D)** Endogenous CD4^+^FOXP3^+^ Tregs frequency in the heart and spleen **(E)** of control no-transfer (gray) and TCR-M-transferred (light green) mice. Bars represent mean, error represents SEM and circles illustrate individual samples. Data were acquired from two independent experiments, n=8-12 mice. Statistical analyses in Panels **B** and **C**: One-way ANOVA followed by Tukey’s post-test. *P<0.05 and **P<0.01. Statistics in **D** and **E**: Two-tailed unpaired T test. **P<0.01.

### The infarcted myocardium steers T_H_ polarized TCR-M cells towards regulatory T-cell phenotype

Herein, we report evidence that naive TCR-M cells acquire an iTreg signature poised with suppressive functions in the heart and MedLNs in the context of MI. Some conventional TCR-M cells remain in both sites, though, and the differential roles played by distinct TCR-M subsets with regard to post-MI repair remain unclear. Therefore, we pre-differentiated TCR-M cells towards T_H_1, T_H_17 or Treg phenotypes *in vitro* before transferring them into DO11.10 recipients, one day before MI induction. We opted to use DO11.10 recipients in these experimental sessions to avoid clonal competition between transferred and endogenous cells. Successful enrichment of each T helper cell state of interest was confirmed prior to cell transfer (**Figure 5A**, ∼80% T-bet^+^ IFN-γ^var^ for T_H_1, ∼90% ROR-γt^+^ IL-17^var^ for T_H_17 and ∼43% CD25^+^FOXP3^+^ for Tregs). Flow cytometry analysis revealed a stark accumulation of T_H_1-polarized TCR-M cells in the infarcted myocardium (**Figure 5B**). The T_H_1-transferred cells found in the hearts of infarcted recipients largely retained the pro-inflammatory phenotypic signature (T-bet^+^) on day 5 post MI (**Figure 5B**). The T_H_17-polarized TCR-Ms found in the infarcted hearts showed reduced ROR-γt expression compared to pre-transfer levels but still presented higher expression than the spleen counterparts (**Figure 5C**). In contrast to T_H_1 polarized group, 17.1 % (± 4.4%) of T_H_17-polarized TCR-M cells found in the infarcted hearts acquired FOXP3 expression (in contrast to 2% pre-transfer levels). The splenic TCR-M cells retained their T_H_17 phenotype similar to pre-transfer levels, reinforcing the notion of myocardial conversion (**Figure 5C**). In line with those observations, Treg-enriched TCR-M cells kept steady FOXP3 expression in all sites, at levels similar to those obtained pre-transfer (**Figure 5D**). We found no evidence that Tregs lose FOXP3 expression in the acutely injured myocardium. Altogether, these experiments provide functional evidence supporting the idea that the infarcted myocardium signals to myosin-specific T helper cells to shape their phenotypic plasticity towards a regulatory phenotype marked by FOXP3 expression.

**Figure 5.**
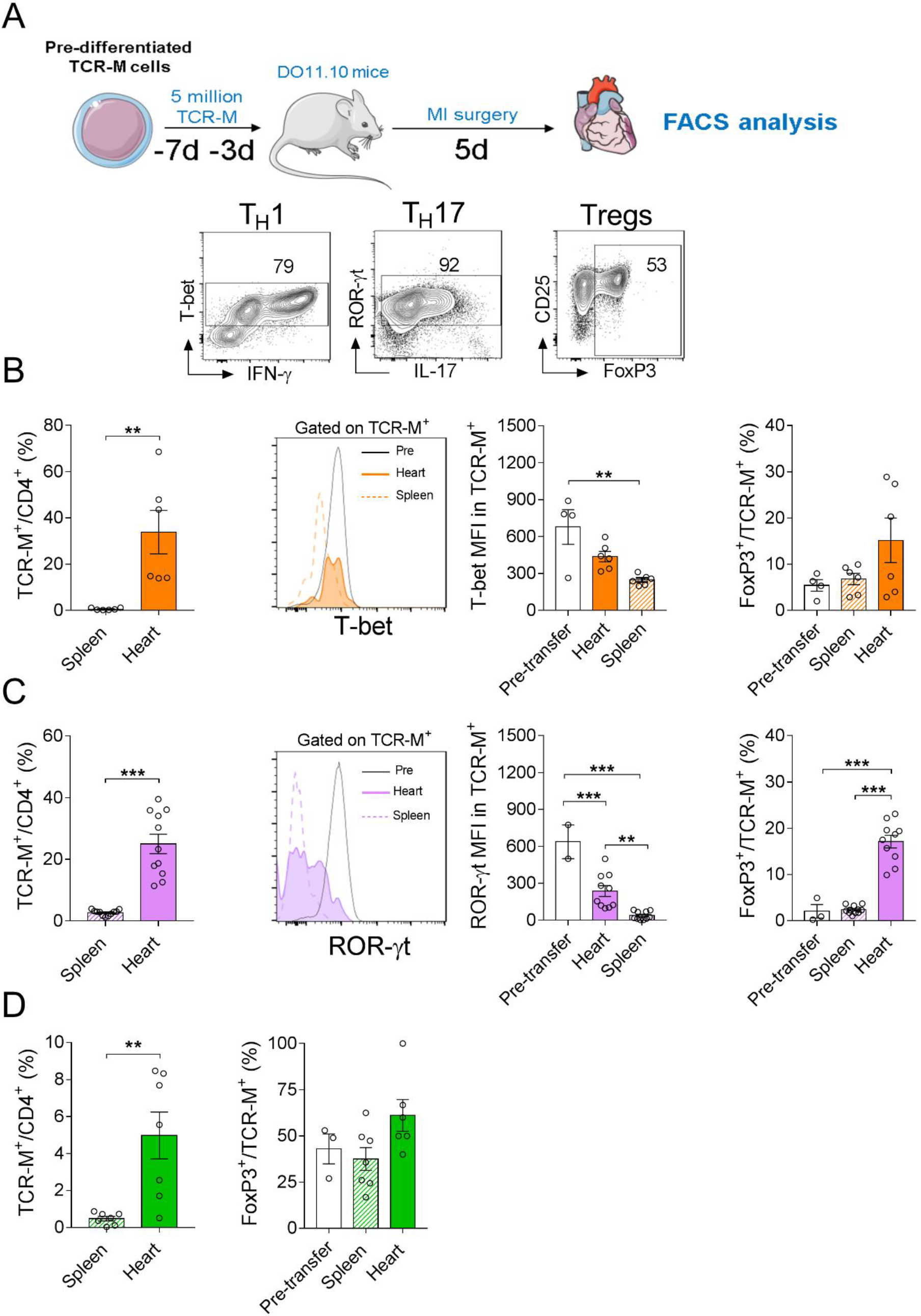
The infarcted myocardium steers polarized TCR-M cells towards a regulatory phenotype. (**A**) TCR-M cells pre-differentiated towards T_H_1, T_H_17 and Tregs were inject into DO11.10 hosts one day prior to MI surgery. FACS analysis of heart and spleen tissue was performed 5 days post-MI. Contour-plots illustrate the expression of prototypic TFs and cytokines for each polarization state at pre-transfer level. (**B**) Frequency of T_H_1 TCR-M cells among CD4^+^ T-cells in the spleen and heart after MI. Overlaid histogram and graph depicts T-bet expression in the transferred TCR-M cells found in spleens (dashed lines) and hearts (filled histogram) of recipients. Open bars represent pre-transfer MFI. Frequency of FOXP3^+^ TCR-M^+^ cells in T_H_1 polarized cells at pre-transfer or analyzed in the heart and spleen tissue 5d post-MI. (**C**) Frequency of T_H_17 TCR-M cells among heart and spleen CD4^+^ T-cells. Histogram illustrates ROR-γt expression in T_H_17 polarized TCR-M cells from pre-transfer or heart and spleen isolated cells. Frequency of FOXP3^+^ T_H_17 TCR-Ms at pre-transfer or obtained from spleen and heart at 5d post-MI. (**D**) Frequency of Treg polarized TCR-M cells among CD4^+^ T-cells in the heart and spleen at 5d post-MI. Frequency of FOXP3^+^ in Treg polarized TCR-M cells at pre-transfer or isolated from heart and spleen tissue. Bar graphs depict the mean, SEM and the distribution of individual samples (4-11 mice per group). Statistical analysis in panels **B, C and D** left panel Two-tailed unpaired T test. ***P<0.001. Statistical analysis in **B, C and D right panel**: one-way ANOVA followed by Tukey’s *post hoc*. *P<0.05, **P<0.01 and ***P<0.001.

### Dissecting the differential contribution of distinct TCR-M phenotypic states to post-MI responses

Next, we sought to dissect how each distinct TCR-M phenotypic state can influence myocardial inflammatory response and cardiac function after MI. Along with the experimental groups reported in **Figure 5**, infarcted DO11.10 mice that did not receive adoptive T-cell transfer were used as controls. Post-MI responses were monitored by echocardiography and flow cytometry performed on day 5 after MI. Flow cytometry analysis of digested myocardial scars revealed that TCR-M T_H_1-transferred mice exhibit an increased recruitment of pro-inflammatory Ly6C^hi^ monocytes (**Figure 6A**), whereas adoptive transfer of TCR-M Tregs resulted in an overall decrease in myocardial leukocyte infiltrate (**Figure 6A**). Gene expression analysis of pro-inflammatory transcripts (*Il1b* and *Tnf*) further confirmed that adoptive transfer of T_H_1 and T_H_17 polarized TCR-M cells fueled cardiac inflammation, in contrast to Treg-transferred group that favored the expression of the cardiomyocyte-related transcript *Myh7* (**Figure 6B)**.These observations indicate that, despite being present at rather low frequencies in the injured myocardium, antigen-specific T-cells can shape the local inflammatory milieu, either fueling or suppressing *in situ* responses mirroring their phenotypic states.

**Figure 6.**
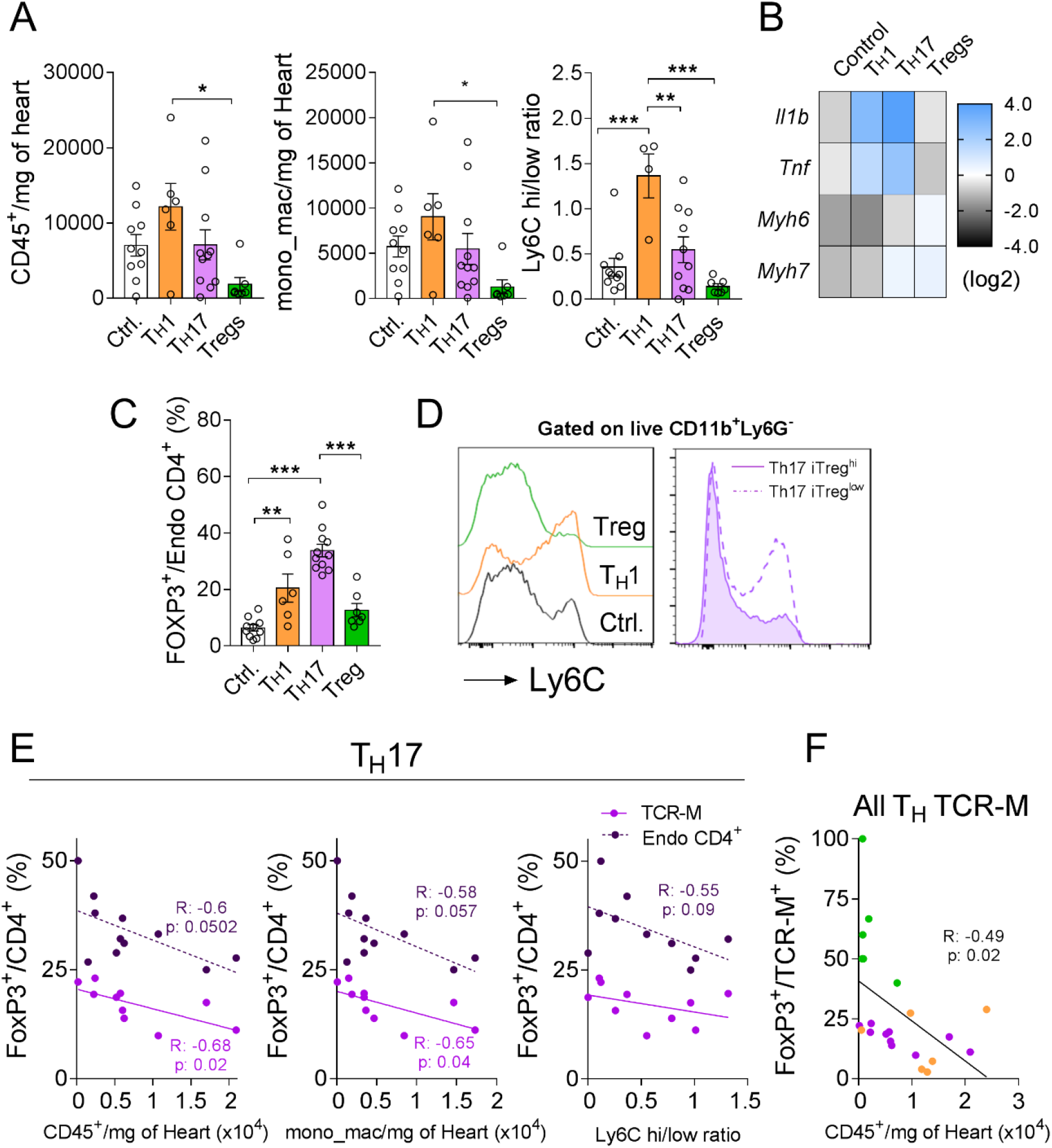
Differential regulation of myocardial infarction inflammation by TCR-M cells with distinct phenotypes. **(A)** Numbers of leukocytes, monocytes/macrophages; and ratio of Ly6CHi/Low monocytes in the hearts of T_H_ TCR-M-transferred mice. (**B**) Normalized gene expression of pro-inflammatory and cardiomyocyte related transcripts in the heart of TCR-M T_H_ transferred mice. (**C**) Frequency of endogenous cardiac Tregs in distinct TCR-M T_H_ transferred mice. (**D**) Overlaid histogram (left panel) illustrates the frequency of Ly6C^Hi^ monocytes in the heart of control (Ctrl.), T_H_1 and Treg transferred mice. Overlaid histogram (right panel) indicates the number of Ly6C^Hi^ monocytes in T_H_17 transferred mice with high or low endogenous Treg numbers. (**E**) Correlation of induced TCR-M Tregs (light purple) and endogenous Tregs (dark purple) against heart CD45^+^ counts, monocyte/macrophage counts and Ly6CHi/low ratio in T_H_17 transferred mice. (**F**) Correlation of induced TCR-M Tregs in T_H_1 (orange), T_H_17 (purple) and Treg (green) transferred mice against respective heart CD45^+^ counts. Bars represent mean, error represents SEM and symbols illustrate individual samples. N=6-11 mice per group. Open bars represent no TCR-M-transferred control mice. Statistical analysis in **A**: one-way ANOVA followed by Tukey’s *post hoc*. *P<0.05, **P<0.01, ***P<0.001. Statistical analysis in **E** and **F**: Pearson correlation, respective R and p values are indicated in the figure.

Surprisingly, the adoptive transfer of T_H_17 TCR-M cells (and to a lesser extent of T_H_1 TCR-M cells) promoted an expansion of the recipients’ endogenous FOXP3^+^ Treg population (**Figure 6C**). These findings reveal complex crosstalk among different T-cell subsets in the healing myocardium. In light of these observations, we decided to explore a possible relationship between the fate of T_H_17-transferred cells and the in situ inflammatory response. As shown in **Figure 6D-E**, we found that the levels of myocardial leukocyte and myeloid cell numbers negatively correlated with the T_H_17-to-Treg conversion. Likewise, the myocardial inflammatory cell infiltration also negatively correlated with the TCR-M-induced expansion of the endogenous Treg compartment. Moreover, a lower Ly6C^hi^ pro-inflammatory monocyte frequency correlated to higher endogenous Treg counts in this experimental group (**Figure 6E**). Taken together, these findings indicate that in vivo T_H_17-to-Treg conversion efficiently blocked myocardial inflammation similar to the effects observed for Tregs that had been pre-differentiated in vitro (**Figure 6F**).

Echocardiography analysis performed on control and T_H_-transferred TCR-M mice, on day 5 after MI, revealed that while TCR-M cells had no impact on infarct size or survival (**Supplementary table 1**), Treg-transferred mice showed better-preserved end systolic and diastolic areas (ESA, EDA) (**Supplementary table 1**). Mice transferred with T_H_17-polarised TCR-M cells also showed improved cardiac function (**Supplementary table 1**), but it remains unclear whether this is related to their partial Treg conversion or to other yet unknown mechanisms. These results demonstrate complex and dynamic crosstalk between different T helper cell subsets in the healing myocardium, which ultimately contributes to more effective tissue repair.

## Discussion

In the present study, we performed deep phenotyping of antigen-specific post-MI responses and found that myosin-specific T helper cells develop an induced regulatory signature in the heart and MedLNs of infarcted mice, with cells mainly differentiating towards either an “*effector Treg*” or “*suppressor*” state, associated with fibrotic repair and local immune response regulation. Functional *in vivo* experiments performed using two different transgenic TCR models (MYHCA- and OVA-specific) and adoptive transfer of pre-differentiated TCR-M cells confirmed that the infarcted myocardial milieu directs the antigen-specific T helper cells to develop a regulatory phenotype and to suppress local inflammatory responses. Moreover, in contrast to other chronic myocardial diseases^15, 16, 33^, no evidence for loss of FOXP3 expression (i.e., ex-Treg) was observed in the acute phase of post-MI responses, further confirming that the cardio-immune crosstalk at this early stage favors salutary adaptive immune mechanisms. Finally, adoptive cell transfer experiments using TCR-M cells pre-differentiated towards defined phenotypic signatures revealed complex crosstalk, between conventional (FOXP3^-^) and regulatory (FOXP3^+^) cells in the infarcted milieu, that ultimately favored myocardial healing.

In a previous study, we identified a peptide from the cardiac myosin heavy alpha chain (MYHCA_614-629_) as a dominant myocardial antigen triggering cardioprotective CD4^+^ T-cell responses in the MI context^7^, but the precise phenotype of these antigen-specific T-cells in post-MI responses has not been fully elucidated. When exposed to inflammatory conditions, FOXP3-expressing Tregs can exhibit an unstable phenotype and express pro-inflammatory cytokines^34^. For instance, Tregs have been shown to lose their suppressive function, acquire a T_H_1/T_H_17 phenotype and contribute to ischemic heart failure and atherosclerosis progression^15, 16^. We therefore combined deep phenotyping approaches (scRNA/ scTCRseq) with transgenic TCR models of adoptive cell transfer to scrutinize in detail how MI affects the antigen-specific T-cell differentiation.

Single-cell-level transcriptomic profiling of the myocardial-MedLN axis revealed ten CD4^+^ T-cell clusters clearly distinguished from one another by conventional/regulatory phenotype and activation state. Interestingly, TCR-M cells showed a signature compatible with peripherally induced Tregs, marked by the expression of transcripts like *Izumo1r, Tgfb1, Cd200* and *Lag3*, among others^23, 24^. In addition, TCR-M cells lacked a gene expression signature, suggesting conventional T_H_17 polarization. A closer examination of the naive TCR-M cells activated during MI uncovered that they differentiate into an early activation state and then branch into either *effector Treg* or *suppressor* clusters. Notably, the inducible Treg signature developed alongside this differentiation trajectory. The *effector Treg* cells were predominant in infarcted hearts at day 5 and expressed high levels of transcription factor (TF) *Myc*^*35*^, antigen-specific stimulation markers (*Nr4a1* and *Tnfrsf9*), *Mif* (macrophage migration inhibitory factor) and *Tgfb1* (Transforming growth factor beta 1), which have been shown to influence early myocardial repair, cardiomyocyte metabolism, pro-fibrotic responses and immune responses^36-40^. In contrast, the *suppressor* TCR-M cells were characterized by the expression of several immune inhibitory receptors (e.g. *Pdcd1, Ctla4* and *Tigit*)^41^. This cluster also included a mixed gene expression signature compatible with follicular T helper cells (e.g. *Il21, Cxcr5* and *Il4*), but their potential impacts on myocardial antibody production were not addressed in this study. Functional adoptive transfer experiments confirmed the suppressor/regulatory phenotypes of transferred TCR-M cells.

Besides advancing our understanding of post-MI T-cell responses, single-cell-level transcriptomic profiling of heart-specific T-cells purified from sham-operated controls also sheds light on how peripheral mechanisms actively maintain immunological tolerance to cardiac antigens during steady-state conditions. Unlike most known autoantigens, the prototypical cardiac antigen MYHCA is not expressed in the thymus, where central mechanisms of dominant immunological tolerance take place^42^. It is therefore remarkable that even in the absence of myocardial injury, TCR-M cells still show signs of early activation and Treg conversion, though to a lesser extent than in the context of infarction. These observations reinforce that constitutive presentation of cardiac autoantigens occurs in the steady state^43, 44^ and that immunological tolerance to myocardial antigens is actively maintained by peripheral mechanisms that remain poorly understood. That the healthy myocardium expresses higher baseline levels of several immune inhibitory receptors than most other tissues^45^ may also contribute to peripheral tolerance mechanisms, though this requires further investigation. In a recent study, our co-authors reported that TCR-M cells can cross-react with a bacterial antigen encountered in the colonic environment, leading to the development of pathogenic heart-directed T_H_17 responses^46^. However, similar peptide mimicry mechanisms, which develop over the course of several weeks, are unlikely to play a role in the phenomena we observed during acute adoptive cell transfer experiments (days 5-7 post MI).

Though most of the TCR-M cells activated in the MI context acquire a Treg signature, infarcted hearts also contained conventional FOXP3^-^ TCR-M cells. These distinct T-cell subsets’ specific contributions to post-MI inflammation and repair have not yet been dissected. Unexpectedly, our findings revealed that adoptive transfer of conventional TCR-M cells primed toward a pro-inflammatory phenotype also expanded the recipients’ endogenous Treg compartment, revealing complex cooperation mechanisms between these often-antagonistic states. The mechanisms underlying this bystander, T_H_1/T_H_17 induced rise in endogenous Tregs remain elusive. Still, it is plausible to assume that the IL-2 secreted by the transferred pre-activated TCR-M cells might contribute to this phenomenon, in parallel to the Treg expansion instigated by low IL-2 treatment^47, 48^.

Adoptive transfer of T_H_1-predifferentiated TCR-Ms led to increased inflammatory levels in the heart, whereas FOXP3^+^ Tregs blunted the myocardial leukocyte infiltration. The Treg-mediated anti-inflammatory effects were seen both o animals that received Tregs pre-differentiated in vitro and in animals showing high T_H_17-to-Treg conversion in vivo. These findings confirm that antigen-specific T-cells help shape myocardial inflammatory responses and can regulate myocardial inflammatory responses despite being present at low frequencies only. These findings are in line with previous reports indicating myocardial Tregs have salutary effects in the context of MI ^3, 5, 7, 25, 49^, though this is the first experimental evidence using Tregs with defined antigen specificity. Surprisingly, our data showed both T_H_17, and Treg TCR-Ms led to better-preserved EDA and ESA indices on day 5 post MI, when compared to no-transfer infarcted controls. However, it remains unclear whether the protection observed in the T_H_17 transferred group is dependent on Treg conversion or on other unknown mechanisms. It is important to stress that post-MI inflammation is a Janus-faced mechanism and, while overshooting inflammation is obviously detrimental, effective *in situ* inflammation is also vital for proper healing and development of functional scar ^50, 51^.

Taken together, in the present study we dissected how the myocardial milieu steers CD4^+^ T-cell responses towards a regulatory phenotype through signaling molecules and cell activation states associated with effector and suppressor phenotypes. Our findings also provide functional evidence supporting salutary cardio-(auto)immune crosstalk during the acute phase of post-MI repair, maintained via mechanisms of peripheral immunological tolerance and complex cooperation between pro-inflammatory and regulatory T helper cells.

## Acknowledgements

The authors greatly appreciate the skillful technical assistance of Elena Vogel and Andrea Leupold. They thank Dr. Max Rieckmann and Prof Encarnita Mariotti-Ferrandiz for the insightful comments and suggestions.

## Author contributions

VS and PPR analyzed the human myocardial tissue. LP performed mouse echocardiography measurements and analyses. MD, EW, LP, PA-L and GCR conducted experiments and analyzed data (FACS, cell sorting, immunophenotyping). PA and AES conducted the single-cell RNA sequencing experiments. MD, DEA and GCR analyzed the single cell RNA sequencing data. MD, EW, DEA, UH, SF and GCR made substantial contributions to the conception and design of the present work. BL generated and provided the TCR-M mice and contributed to data interpretation. All-coauthors contributed to the manuscript preparation.

## References

1. Swirski FK, Nahrendorf M. Cardioimmunology: The immune system in cardiac homeostasis and disease. Nat Rev Immunol. 2018;18:733–744

2. Ramos GC, van den Berg A, Nunes-Silva V, Weirather J, Peters L, Burkard M, Friedrich M, Pinnecker J, Abesser M, Heinze KG, Schuh K, Beyersdorf N, Kerkau T, Demengeot J, Frantz S, Hofmann U. Myocardial aging as a t-cell-mediated phenomenon. Proc Natl Acad Sci U S A. 2017;114:E2420–E2429

3. Hofmann U, Beyersdorf N, Weirather J, Podolskaya A, Bauersachs J, Ertl G, Kerkau T, Frantz S. Activation of cd4+ t lymphocytes improves wound healing and survival after experimental myocardial infarction in mice. Circulation. 2012;125:1652–1663

4. D’Alessio FR, Kurzhagen JT, Rabb H. Reparative t lymphocytes in organ injury. J Clin Invest. 2019;129:2608–2618

5. Weirather J, Hofmann UD, Beyersdorf N, Ramos GC, Vogel B, Frey A, Ertl G, Kerkau T, Frantz S. Foxp3+ cd4+ t cells improve healing after myocardial infarction by modulating monocyte/macrophage differentiation. Circ Res. 2014;115:55–67

6. Nosbaum A, Prevel N, Truong HA, Mehta P, Ettinger M, Scharschmidt TC, Ali NH, Pauli ML, Abbas AK, Rosenblum MD. Cutting edge: Regulatory t cells facilitate cutaneous wound healing. J Immunol. 2016;196:2010–2014

7. Rieckmann M, Delgobo M, Gaal C, Buchner L, Steinau P, Reshef D, Gil-Cruz C, Horst ENT, Kircher M, Reiter T, Heinze KG, Niessen HW, Krijnen PA, van der Laan AM, Piek JJ, Koch C, Wester HJ, Lapa C, Bauer WR, Ludewig B, Friedman N, Frantz S, Hofmann U, Ramos GC. Myocardial infarction triggers cardioprotective antigen-specific t helper cell responses. J Clin Invest. 2019;129:4922–4936

8. Hammer A, Sulzgruber P, Koller L, Kazem N, Hofer F, Richter B, Blum S, Hulsmann M, Wojta J, Niessner A. The prognostic impact of circulating regulatory t lymphocytes on mortality in patients with ischemic heart failure with reduced ejection fraction. Mediators Inflamm. 2020;2020:6079713

9. Nevers T, Salvador AM, Grodecki-Pena A, Knapp A, Velazquez F, Aronovitz M, Kapur NK, Karas RH, Blanton RM, Alcaide P. Left ventricular t-cell recruitment contributes to the pathogenesis of heart failure. Circ Heart Fail. 2015;8:776–787

10. Delgobo M, Frantz S. Heart failure in cancer: Role of checkpoint inhibitors. Journal of Thoracic Disease. 2018:S4323-S4334

11. Johnson DB, Balko JM, Compton ML, Chalkias S, Gorham J, Xu Y, Hicks M, Puzanov I, Alexander MR, Bloomer TL, Becker JR, Slosky DA, Phillips EJ, Pilkinton MA, Craig-Owens L, Kola N, Plautz G, Reshef DS, Deutsch JS, Deering RP, Olenchock BA, Lichtman AH, Roden DM, Seidman CE, Koralnik IJ, Seidman JG, Hoffman RD, Taube JM, Diaz LA, Jr., Anders RA, Sosman JA, Moslehi JJ. Fulminant myocarditis with combination immune checkpoint blockade. N Engl J Med. 2016;375:1749–1755

12. Sakaguchi S, Vignali DA, Rudensky AY, Niec RE, Waldmann H. The plasticity and stability of regulatory t cells. Nat Rev Immunol. 2013;13:461–467

13. Burzyn D, Kuswanto W, Kolodin D, Shadrach JL, Cerletti M, Jang Y, Sefik E, Tan TG, Wagers AJ, Benoist C, Mathis D. A special population of regulatory t cells potentiates muscle repair. Cell. 2013;155:1282–1295

14. Arpaia N, Green JA, Moltedo B, Arvey A, Hemmers S, Yuan S, Treuting PM, Rudensky AY. A distinct function of regulatory t cells in tissue protection. Cell. 2015;162:1078–1089

15. Bansal SS, Ismahil MA, Goel M, Zhou G, Rokosh G, Hamid T, Prabhu SD. Dysfunctional and proinflammatory regulatory t-lymphocytes are essential for adverse cardiac remodeling in ischemic cardiomyopathy. Circulation. 2019;139:206–221

16. Wolf D, Gerhardt T, Winkels H, Michel NA, Pramod AB, Ghosheh Y, Brunel S, Buscher K, Miller J, McArdle S, Baas L, Kobiyama K, Vassallo M, Ehinger E, Dileepan T, Ali A, Schell M, Mikulski Z, Sidler D, Kimura T, Sheng X, Horstmann H, Hansen S, Mitre LS, Stachon P, Hilgendorf I, Gaddis DE, Hedrick C, Benedict CA, Peters B, Zirlik A, Sette A, Ley K. Pathogenic autoimmunity in atherosclerosis evolves from initially protective apolipoprotein b100-reactive cd4(+) t-regulatory cells. Circulation. 2020;142:1279–1293

17. Li J, Tan J, Martino MM, Lui KO. Regulatory t-cells: Potential regulator of tissue repair and regeneration. Front Immunol. 2018;9:585

18. Nindl V, Maier R, Ratering D, De Giuli R, Zust R, Thiel V, Scandella E, Di Padova F, Kopf M, Rudin M, Rulicke T, Ludewig B. Cooperation of th1 and th17 cells determines transition from autoimmune myocarditis to dilated cardiomyopathy. Eur J Immunol. 2012;42:2311–2321

19. Keppner L, Heinrichs M, Rieckmann M, Demengeot J, Frantz S, Hofmann U, Ramos G. Antibodies aggravate the development of ischemic heart failure. Am J Physiol Heart Circ Physiol. 2018;315:H1358–H1367

20. Lindsey ML, Bolli R, Canty JM, Jr., Du XJ, Frangogiannis NG, Frantz S, Gourdie RG, Holmes JW, Jones SP, Kloner RA, Lefer DJ, Liao R, Murphy E, Ping P, Przyklenk K, Recchia FA, Schwartz Longacre L, Ripplinger CM, Van Eyk JE, Heusch G. Guidelines for experimental models of myocardial ischemia and infarction. Am J Physiol Heart Circ Physiol. 2018;314:H812–H838

21. rodents Fwgorogfhmo, rabbits, Mahler Convenor M, Berard M, Feinstein R, Gallagher A, Illgen-Wilcke B, Pritchett-Corning K, Raspa M. Felasa recommendations for the health monitoring of mouse, rat, hamster, guinea pig and rabbit colonies in breeding and experimental units. Lab Anim. 2014;48:178–192

22. Zarak-Crnkovic M, Kania G, Jazwa-Kusior A, Czepiel M, Wijnen WJ, Czyz J, Muller-Edenborn B, Vdovenko D, Lindner D, Gil-Cruz C, Bachmann M, Westermann D, Ludewig B, Distler O, Luscher TF, Klingel K, Eriksson U, Blyszczuk P. Heart non-specific effector cd4(+) t cells protect from postinflammatory fibrosis and cardiac dysfunction in experimental autoimmune myocarditis. Basic Res Cardiol. 2019;115:6

23. Zemmour D, Zilionis R, Kiner E, Klein AM, Mathis D, Benoist C. Single-cell gene expression reveals a landscape of regulatory t cell phenotypes shaped by the tcr. Nat Immunol. 2018;19:291–301

24. Stubbington MJ, Mahata B, Svensson V, Deonarine A, Nissen JK, Betz AG, Teichmann SA. An atlas of mouse cd4(+) t cell transcriptomes. Biol Direct. 2015;10:14

25. Xia N, Lu Y, Gu M, Li N, Liu M, Jiao J, Zhu Z, Li J, Li D, Tang T, Lv B, Nie S, Zhang M, Liao M, Liao Y, Yang X, Cheng X. A unique population of regulatory t cells in heart potentiates cardiac protection from myocardial infarction. Circulation. 2020;142:1956–1973

26. Cano-Gamez E, Soskic B, Roumeliotis TI, So E, Smyth DJ, Baldrighi M, Wille D, Nakic N, Esparza-Gordillo J, Larminie CGC, Bronson PG, Tough DF, Rowan WC, Choudhary JS, Trynka G. Single-cell transcriptomics identifies an effectorness gradient shaping the response of cd4(+) t cells to cytokines. Nat Commun. 2020;11:1801

27. Lv H, Havari E, Pinto S, Gottumukkala RV, Cornivelli L, Raddassi K, Matsui T, Rosenzweig A, Bronson RT, Smith R, Fletcher AL, Turley SJ, Wucherpfennig K, Kyewski B, Lipes MA. Impaired thymic tolerance to alpha-myosin directs autoimmunity to the heart in mice and humans. J Clin Invest. 2011;121:1561–1573

28. Kim HJ, Barnitz RA, Kreslavsky T, Brown FD, Moffett H, Lemieux ME, Kaygusuz Y, Meissner T, Holderried TA, Chan S, Kastner P, Haining WN, Cantor H. Stable inhibitory activity of regulatory t cells requires the transcription factor helios. Science. 2015;350:334–339

29. Xu C, Fu Y, Liu S, Trittipo J, Lu X, Qi R, Du H, Yan C, Zhang C, Wan J, Kaplan MH, Yang K. Batf regulates t regulatory cell functional specification and fitness of triglyceride metabolism in restraining allergic responses. J Immunol. 2021;206:2088–2100

30. Miragaia RJ, Gomes T, Chomka A, Jardine L, Riedel A, Hegazy AN, Whibley N, Tucci A, Chen X, Lindeman I, Emerton G, Krausgruber T, Shields J, Haniffa M, Powrie F, Teichmann SA. Single-cell transcriptomics of regulatory t cells reveals trajectories of tissue adaptation. Immunity. 2019;50:493–504 e497

31. McFadden C, Morgan R, Rahangdale S, Green D, Yamasaki H, Center D, Cruikshank W. Preferential migration of t regulatory cells induced by il-16. J Immunol. 2007;179:6439–6445

32. Farbehi N, Patrick R, Dorison A, Xaymardan M, Janbandhu V, Wystub-Lis K, Ho JW, Nordon RE, Harvey RP. Single-cell expression profiling reveals dynamic flux of cardiac stromal, vascular and immune cells in health and injury. Elife. 2019;8

33. Bansal SS, Ismahil MA, Goel M, Patel B, Hamid T, Rokosh G, Prabhu SD. Activated t lymphocytes are essential drivers of pathological remodeling in ischemic heart failure. Circ Heart Fail. 2017;10:e003688

34. Sharma A, Rudra D. Emerging functions of regulatory t cells in tissue homeostasis. Front Immunol. 2018;9:883

35. Saravia J, Zeng H, Dhungana Y, Bastardo Blanco D, Nguyen TM, Chapman NM, Wang Y, Kanneganti A, Liu S, Raynor JL, Vogel P, Neale G, Carmeliet P, Chi H. Homeostasis and transitional activation of regulatory t cells require c-myc. Sci Adv. 2020;6:eaaw6443

36. Miller EJ, Li J, Leng L, McDonald C, Atsumi T, Bucala R, Young LH. Macrophage migration inhibitory factor stimulates amp-activated protein kinase in the ischaemic heart. Nature. 2008;451:578–582

37. Qi D, Atsina K, Qu L, Hu X, Wu X, Xu B, Piecychna M, Leng L, Fingerle-Rowson G, Zhang J, Bucala R, Young LH. The vestigial enzyme d-dopachrome tautomerase protects the heart against ischemic injury. J Clin Invest. 2014;124:3540–3550

38. Ikeuchi M, Tsutsui H, Shiomi T, Matsusaka H, Matsushima S, Wen J, Kubota T, Takeshita A. Inhibition of tgf-beta signaling exacerbates early cardiac dysfunction but prevents late remodeling after infarction. Cardiovasc Res. 2004;64:526–535

39. Kuwahara F, Kai H, Tokuda K, Kai M, Takeshita A, Egashira K, Imaizumi T. Transforming growth factor-beta function blocking prevents myocardial fibrosis and diastolic dysfunction in pressure-overloaded rats. Circulation. 2002;106:130–135

40. Sanjabi S, Oh SA, Li MO. Regulation of the immune response by tgf-beta: From conception to autoimmunity and infection. Cold Spring Harb Perspect Biol. 2017;9

41. Rumpret M, Drylewicz J, Ackermans LJE, Borghans JAM, Medzhitov R, Meyaard L. Functional categories of immune inhibitory receptors. Nat Rev Immunol. 2020;20:771–780

42. Coutinho A. The le douarin phenomenon: A shift in the paradigm of developmental self-tolerance. Int J Dev Biol. 2005;49:131–136

43. Smith SC, Allen PM. Expression of myosin-class ii major histocompatibility complexes in the normal myocardium occurs before induction of autoimmune myocarditis. Proc Natl Acad Sci U S A. 1992;89:9131–9135

44. Van der Borght K, Scott CL, Nindl V, Bouche A, Martens L, Sichien D, Van Moorleghem J, Vanheerswynghels M, De Prijck S, Saeys Y, Ludewig B, Gillebert T, Guilliams M, Carmeliet P, Lambrecht BN. Myocardial infarction primes autoreactive t cells through activation of dendritic cells. Cell Rep. 2017;18:3005–3017

45. Ashour D, Delgobo M, Frantz S, Ramos GC. Coping with sterile inflammation: Between risk and necessity. Cardiovasc Res. 2021;117:e84–e87

46. Gil-Cruz C, Perez-Shibayama C, De Martin A, Ronchi F, van der Borght K, Niederer R, Onder L, Lutge M, Novkovic M, Nindl V, Ramos G, Arnoldini M, Slack EMC, Boivin-Jahns V, Jahns R, Wyss M, Mooser C, Lambrecht BN, Maeder MT, Rickli H, Flatz L, Eriksson U, Geuking MB, McCoy KD, Ludewig B. Microbiota-derived peptide mimics drive lethal inflammatory cardiomyopathy. Science. 2019;366:881–886

47. Klatzmann D, Abbas AK. The promise of low-dose interleukin-2 therapy for autoimmune and inflammatory diseases. Nat Rev Immunol. 2015;15:283–294

48. Zhao TX, Kostapanos M, Griffiths C, Arbon EL, Hubsch A, Kaloyirou F, Helmy J, Hoole SP, Rudd JHF, Wood G, Burling K, Bond S, Cheriyan J, Mallat Z. Low-dose interleukin-2 in patients with stable ischaemic heart disease and acute coronary syndromes (lilacs): Protocol and study rationale for a randomised, double-blind, placebo-controlled, phase i/ii clinical trial. BMJ open. 2018;8:e022452

49. Saxena A, Dobaczewski M, Rai V, Haque Z, Chen W, Li N, Frangogiannis NG. Regulatory t cells are recruited in the infarcted mouse myocardium and may modulate fibroblast phenotype and function. Am J Physiol Heart Circ Physiol. 2014;307:H1233–1242

50. Frantz S, Hofmann U, Fraccarollo D, Schafer A, Kranepuhl S, Hagedorn I, Nieswandt B, Nahrendorf M, Wagner H, Bayer B, Pachel C, Schon MP, Kneitz S, Bobinger T, Weidemann F, Ertl G, Bauersachs J. Monocytes/macrophages prevent healing defects and left ventricular thrombus formation after myocardial infarction. FASEB J. 2013;27:871–881

51. Vagnozzi RJ, Maillet M, Sargent MA, Khalil H, Johansen AKZ, Schwanekamp JA, York AJ, Huang V, Nahrendorf M, Sadayappan S, Molkentin JD. An acute immune response underlies the benefit of cardiac stem cell therapy. Nature. 2020;577:405–409

