## Supplemental material for "Rapid differentiation of regulatory CD4^+^ T cells in the infarcted myocardium blunts in situ inflammation"

**Rapid differentiation of regulatory CD4[+] T cells in the infarcted myocardium blunts in situ inflammation and preserves cardiac function**

Authors:

Murilo Delgobo[1, 2], Emil Weiß[1, 2], Diyaa ElDin Ashour,[1, 2] Lisa Popiolkowski[1, 2], Panagiota Arampatzi[3], Verena Stangl[4], Paula Arias-Loza[5], Peter P. Rainer[4, 6], Antoine-Emmanuel Saliba[7], Burkhard Ludewig[8], Ulrich Hofmann[1, 2], Stefan Frantz[1, 2], and Gustavo Campos Ramos[1, 2, §]

Supplemental material

Supplemental methods

Supplemental Figures 1-5

Supplementary table 1

Supplemental references

### Supplemental Methods

*Animals:* The mice were maintained in individually ventilated cages under specific pathogen free (SPF) conditions with a 12-hour light/12-hour dark cycle and standard diet provided ad libitum, and enrolled in experimental procedures at the age of 8-12 weeks. Thy1.2 WT male BALB/c mice were commercially acquired from Charles River (Sulzfeld, Germany). TCR-M mice expressing a transgenic TCR specific for a class-II-restricted cardiac myosin peptide ($MYHCA_{614-629}$) were bred in the housing facility at the University Clinic Würzburg after being provided by Prof. Burkhard Ludewig[1]. TCR-M mice were bred with BALB/c Thy1.1 mice to express the congenic marker Thy1.1. Thy1.2 DO11.10 expressing a transgenic TCR against

$OVA_{323-339}$ peptide were bred in our local housing facility at the University Clinic Würzburg. All mouse strains used in this study shared the same genetic background (BALB/c). All in vivo procedures have been approved by the local authorities (*Regierung von Unterfranken*) and in conform with the Federation for Laboratory Animal Science Associations (FELASA) guidelines.

*Experimental myocardial infarction (EMI):* Surgery was performed in aseptic conditions and myocardial infarction was induced by the permanent ligation of the left coronary artery (LAD) as previously described [2]. Mice were anesthetized with isoflurane (Forene, Abbott, Wiesbaden, Germany), intubated for mechanical ventilation and subjected to thoracotomy. Mouse skin was shaved, disinfected (Braunoderm, B. Braun, Melseungen, Germany) and an incision was made to access the third intercostal space by blunt preparation and held open by a micro-retractor. After lung displacement using a Ringer lactate-soaked (Baxter, Unterscleißheim, Germany) piece of sponge (Pro.ophta, Lohmann & Rauscher, Rengsdorf, Germany), the LAD of the mice in MI group was permanently ligated with a 6/0 perma-hand silk thread (Ethicon, Nordersteld, Germany). This step was omitted in sham operated mice. The chest cage and skin were then sutured sequentially using a 5/0 and 6/0 Prolene thread, respectively. Isoflurane concentration was reduced to 0.5% and mice were extubated as soon as spontaneous respiration occurred. Body temperature was controlled during the operation, and the mice were subjected to a pain management program (buprenorphine s.c. 0.1 mg/Kg body weight, Buprenovet, Bayer Vital, Leverkusen, Germany) one hour before and for at least 3 days (twice daily) after operation. Endpoint analyses were performed on days 5 and 7 post-operation, preceded by echocardiography in each section.

*Echocardiographic assessments:* Echocardiography measurements were performed with a Vevo 1100 instrument (VisualSonics, Amsterdam, Netherlands) with a MS 400 echo transducer coupled to a 30- MHz probe developed for mice studies. Briefly, mice were maintained under slight isoflurane anesthesia (0.5-1.5% vol/vol O2) on a heated bed (39°C), and images were acquired on the short axis at apex, midpapillary (B and M mode) and basal (only B mode) levels and on the long axis (B and M mode) to represent the infarct border zones. The analyses were performed blinded using the Vevo LAB 3.2.6 software. As inclusion criteria, only mice with infarct size above 20% were used in experiments. The infarct size was measured in the longitudinal axis (B mode) by calculating a quotient from the heart size and the akinetic myocardium (scar tissue)[3].

*Adoptive cell transfer*

*Adoptive transfer of TCR-M cells into MI recipients:* Thy1.1$^+$ myosin-specific CD4$^+$ T-cells were purified from naïve TCR-M mice by magnetic cell sorting and then transferred into Thy1.2$^+$ WT syngenic recipients one day prior to EMI induction. In brief, spleen and lymph nodes from TCR-M mice were aseptically extracted and stored in Hank's balanced salt solution containing 0.5% (wt/vol) bovine serum albumin (HBSS/BSA). A cell suspension was obtained after grinding the lymphoid organs against a 30 µm filter mesh (Miltenyi Biotec, Bergisch-Gladbach, Germany) in HBSS/BSA buffer. Untouched CD4$^+$ T cells were purified by magnetic cell sorting, using the CD4$^+$ T-cell Isolation kit II (Miltenyi Biotec, Bergisch Gladbach, Germany). Prior to injection, TCR-M cells were resuspended in sterile PBS (Biochrom, Berlin, Germany) at a concentration of 2.5x10$^7$ cells/mL and adoptively transferred into recipient mice (5x10$^6$ cells per mice, i.p.). The distribution and phenotype of TCR-M cells in the recipient mice was monitored by flow cytometry.

*Adoptive transfer of DO11.10 cells into MI recipients:* Spleens from DO11.10 mice, containing OVA-specific CD4$^+$ T-cells, were harvested in aseptic conditions and stored in HBSS/BSA buffer. Single-cell suspension was obtained after grinding the spleens against a 30 µm filter mesh (Miltenyii Biotec, Bergisch-Gladbach, Germany) in HBSS/BSA buffer. Cells were washed and resuspended in complete RPMI 1640 media containing 10% FCS at the density of 2x$10^6$ cells/mL in a 24-well plate. For OVA-specific $CD4^+$ T-cell activation, splenocytes were stimulated with 1 µg/mL $OVA_{323-339}$ peptide during 3 days. Unstimulated cells were kept in media with recombinant mouse IL-7 (Peprotech, Hamburg, Germany) at 3.125 ng/mL. After the stimulation period, cells were washed with sterile PBS (Biochrom, Berlin, Germany) and $CD4^+$ T-cells were isolated by negative magnetic selection using the $CD4^+$ T-cell Isolation kit II (Miltenyi Biotec, Bergisch Gladbach, Germany). Before adoptive transfer, activated and resting cells were labeled with distinct cell-tracer dyes. Briefly, activated OVA cells were resuspended in a solution containing 5 µM CFSE (ThermoFisher Scientific, Waltham, MA, USA), while resting OVA cells were resuspended in 5 µM VIO (ThermoFisher Scientific, Waltham, MA, USA), at a density of $10^6$ cells/mL and incubated at 37°C for 20 minutes. Complete RPMI media containing 10% FCS was then added, and cells were maintained for 5 minutes at 37°C. Cell suspensions were then washed and resuspended in sterile PBS. The labeled activated / resting OVA $CD4^+$ T-cells were adoptively transferred into syngenic DO11.10 recipient mice (5x$10^6$ cells, i.p., at 1:1 ratio) one day prior to MI or sham operation. DO11.10 cells were identified in vivo based on respective cell-tracer signal in analyzed cardiac and lymphoid tissues.

*Adoptive transfer of polarized TCR-M cells into MI recipients:* Untouched $CD4^+$ TCR-M cells were separated by magnetic cell sorting as previously described. For $CD4^+$ T-helper polarization, TCR-M cells were resuspended in complete RPMI media supplemented with 10% FCS and seeded at 2x$10^6$ cells/mL in 24-well plate. The following cytokines and neutralizing antibodies were used according to the $T_H$ subset: For $T_H1$ polarization, cells were stimulated with anti CD3/CD28 dynabeads (ThermoFisher Scientific, Waltham, MA, USA) at 1:1 bead to cell ratio, recombinant murine IL-2 500 U/mL and recombinant murine IL-12p70 600 U/mL (Peprotech, Hamburg, Germany) and anti-IL-4 at 5 µg/mL (Biolegend, San Diego, CA, USA, clone 11B11). For $T_H17$ polarization, cells were stimulated with anti CD3/CD28 dynabeads (thermofisher) at 1:1 bead to cell ratio, recombinant murine IL-6 10000 U/mL, recombinant murine IL-1β 8400 U/mL, recombinant murine IL-23 40 U/mL, recombinant human TGF-β 10 U/mL (Peprotech, Hamburg, Germany) and the following neutralizing antibodies: anti-IL-2 at 10 µg/mL (clone JES6-5H4), anti-IL-4 at 5 µg/mL (clone 11B11) and anti-IFN-γ at 10 µg/mL (clone XMG1.2) all obtained from Biolegend®. Cells were kept for three days in stimulation media For Treg expansion, $CD25^+$ cells were enriched and stimulated with anti CD3/CD28 dynabeads (thermofisher) at 2:1 bead to cell ratio and recombinant murine IL-2 2000 U/mL for seven days, with new media addition every 48h. Before adoptive transfer, antiCD3/CD28 beads were removed by incubating wells in a magnetic plate for two minutes, followed by two washing steps with sterile PBS. Cells were resuspend in sterile PBS at $2.5x10^7$ cells/mL density and $5x10^6$ cells were injected i.p. in DO11.10 hosts one day prior to surgery. In each experiment, $T_H$ polarization was confirmed by flow cytometry using the respective antibodies: IFN-γ and T-bet for $T_H1$, IL-17A plus ROR-γt for $T_H17$ and CD25 plus FOXP3 for Tregs. Echocardiography and cardiac leukocyte composition were analyzed five days after experimental MI induction.

*Flow cytometry and cell sorting*

*In Vitro Stimulation Assay for intracellular cytokine detection*

Infarcted mice splenocytes and digested heart tissues were resuspended in complete RPMI media containing 10% FCS, 1% L-Glutamine, 1% Sodium pyruvate, 1% non-essential amino acids, 1%Pen/Strep, 1 µM 2-ME (Gibco – Grovemont Cir, USA) and seeded in U-bottom 96-well plate. Cells were then stimulated for 3 h with a T-cell stimulation cocktail containing Phorbol 12-Myristat 13-Acetat (81 ηM) und Ionomycin (1.34 µM) supplemented with protein transport inhibitors Brefeldin A (10.6 µM) and Monensin (2µM) (eBioscience – San Diego, USA). Afterwards, cells were used in intracellular flow cytometry analysis to detect cytokine expression profile.

*Immunophenotyping*

Infarcted and sham operated mice were sacrificed by cervical dislocation at 5 or 7 days after surgery. Upon euthanasia, mice were perfused with PBS-heparin (50 U/mL) solution to flush coronary circulation and remove blood-borne leukocytes. Heart tissue was enzymatically digested with type II collagenase (1000 IU/mL, Worthington Biochemical Corporation, Lakewood, NJ, USA) for 30 minutes at 37°C. Digested hearts, lymph nodes and spleen samples were filtered through a 30 µm filter mesh (Miltenyi Biotec, Bergisch Gladbach, Germany) in HBSS/BSA buffer. Samples were then washed in PBS and stained with AmCyan Zombie Aqua fixable viability dye (Biolegend, San Diego, CA, USA) for 15 minutes at room temperature. Following a wash step with FACS buffer (PBS containing 1% BSA, 0.1% sodium azide), surface staining was performed in the presence of Fc blocking antibody (anti-CD16/CD32, clone 2.4G2, BD Pharmingen), using the following antibody clones (conjugated with different fluorophores) commercially acquired from Biolegend (San Diego, CA, USA) unless indicated: anti-CD45 (clone 30-F11), anti-TCRβ (clone H57-597), anti-CD4 (clone RM4-5), anti-CD8a (clone 53-6.7), anti-Thy1.1 (clone OX-7), anti-CD11b (clone M1/70), anti-CD25 (clone PC61), anti-B220 (clone RA3-6B2), anti-T-bet (clone 4B10), anti-IFN-γ (clone XMG1.2), anti-IL-17a (clone TC11-18H10.1), anti-LAP (clone TW7-16B4), anti-TNF (clone MP6-XT22). Additionally, the antibodies anti-ROR-γt (clone AFKJS-9), anti-FOXP3 (clone FJK-16s) were obtained from eBioscience and R&D systems respectively. Flow cytometry measurements were performed using the Attune NxT (Thermo Fisher Scientific, Waltham, MA, USA) instrument. Data analysis was performed using Flowjo (Flowjo LLC Ashland, OR, USA). Compensation for spectral overlap was conducted based on single staining controls and the flow cytometry gates were set based on unstained controls.

*Gene expression analysis*

RNA extraction from mouse cardiac tissue (infarcted area) was performed using the tissue RNA isolation kit (RNeasy mini – Qiagen). The RNA concentration and quality were measured in a spectrophotometer and 120 ng was used for cDNA synthesis (iScript – Bio-

Rad). Taqman probes for quantitative PCR were used to measure the expression of pro-inflammatory and cardiomyocyte related transcripts. The following probes were used: *Gapdh* (Mm033002249_g1), *Myh6* (Mm00440359_m1), *Myh7* (Mm01319006_g1), *Tnf* (Mm99999068_m1) and *Il1b* (Mm00434228_m1). mRNA levels were normalized to Gapdh expression and expression level in $T_H$ TCR-M transferred group was normalized to the mean expression value of infarcted non-transferred control group.

*Cell sorting for single cell RNA sequencing*

Digested hearts and MedLNs were obtained from TCR-M transferred mice at 5 and 7 days after sham or MI operation. Cell suspension was obtained as previously described and following surface staining, heart $CD4^+$ T-cells were enriched by magnetic cell sorting, using anti-PE microbeads (Miltenyi Biotec, Bergisch-Gladbach, Germany). Before sorting, samples were separately labeled with an anti-CD45/H2-K hashtag antibody (Biolegend TotalSeqC antibodies, San Diego, CA, USA, 1:150 dilution). The heart and MedLN samples were stained with the following TotalSeq C antibodies: Heart sham 5dpi (CO301), LN sham 5dpi (CO302), Heart MI 5dpi (CO303), LN MI 5dpi (CO304), Heart sham 7dpi (CO305), LN sham 7dpi (CO306), Heart MI 7dpi (CO307) and LN MI 7dpi (CO308). Additionally, all samples were labeled with anti-Thy1.1 (clone OX-7) and anti-CD25 (clone PC61) Cite-Seq antibodies (Biolegend TotalSeqC antibodies, San Diego, CA, USA, 1:100). Sodium azide was omitted in the buffers used for sorting purposes. After staining, heart and MedLN samples were separately pooled and $CD4^+$ T-cells were sorted using a FACS Aria III instrument (BD, Heidelberg, Germany). Following a washing step, all samples were pooled and suspended in 65 µL of PBS/0.04% BSA and loaded in the 10x Genomics Chromium system as one multiplexed sample.

Single cell RNA-sequencing

Chromium controller was used for partitioning single cells into nanolitre-scale Gel Bead-In-Emulsion (GEMs) and Chromium Next GEM Single Cell 5’ v1.1 kits for reverse transcription, cDNA amplification and library construction (10x Genomics, Pleasanton, CA, USA). A

SimpliAmp Thermal cycler was used for amplification and incubation steps (Applied Biosystems, Foster City, CA, USA). Libraries were quantified by a Qubit 3.0 fluorometer (Thermo Fisher Scientific, Waltham, MA, USA) and quality was checked using a 2100 Bioanalyser with high sensitivity DNA kit (Agilent, Santa Clara, CA, USA) in paired-end mode to reach at least 60000 reads per single-cell for gene expression and 6000 reads for the T-cell receptor repertoire, hashtags and Cite-seqs. The Cell Ranger Software suite (10x Genomics, Pleasanton, CA, USA) was used for sequence alignment, barcode processing and sample demultiplexing.

*In silico analysis of single cell sequencing data*

The gene count matrix obtained from Cell Ranger was analyzed using Seurat (version 3.1.4). Cell barcodes detected in gene expression data, hashtag oligo sequences for each group and Cite-seq signal against Thy1.1 antigen were considered during de-multiplexing. For quality control, cells with double hashtag signal or no hashtag detection were excluded. Additionally, cells yielding more than 12.500 unique molecular identifiers (UMI) or more than 5% mitochondrial RNA were also excluded. Cell transcriptome was log-normalized and the top 2,000 most variable genes were obtained using Seurat's default setting. The 20 principal components and a 0.5 resolution were used in dimensionality reduction, resulting in 10 clusters generated by uniform manifold approximation and projection for dimension reduction (UMAP). The function *FindallMarkers* in default settings was then used to identify differently expressed genes between clusters.

The VDJ sequences for each T-cell receptor were added to the Seurat Object using the *combineExpression()* function from scRepertoire R package (https://github.com/ncborcherding/scRepertoire/tree/master/R). The frequency of clonetype occurrence was plotted on a UMAP plot and aligned to CD90.1 TotalSeqC signal to differentiate TCR-M cells from endogenous $CD4^+$ T-cells. The Seurat's function *AddModuleScore* was used to generate gene expression scores related to iTreg, Treg and

$T_H17$ transcriptome. The list of genes describing each cell state in mouse $CD4^+$ T-cell (extended table 1) was normalized based on the expression level of 200 randomly selected co-expressed control genes.

The pseudotime trajectories were built using Monocle3 (version 0.2.3) R package. A Seurat object containing only TCR-M cells was re-clustered and differently expressed genes were obtained using *FindallMarkers* function. The Seurat object was then used to construct a Monocle3 *CellDataSet* (cds) object. Trajectory was plotted using *plot_cells* function and the first node in *Naive1* cluster was set as starting point. The pseudotime values were added back to the Seurat object and represented with a UMAP plot.

The Nichenet analysis was conducted between *effector, suppressor and early/$T_{CM}$* TCR-M cells paired to single cell data containing fibroblasts, macrophages and endothelial cells from infarcted hearts[4]. Following data integration and normalization, the Nichenet R package (https://github.com/saeyslab/nichenetr) was used to build a ligand-target prior model, the ligand receptor network and the weighted integrated networks. In each iteration, effector Tregs, suppressor and early/$T_{CM}$ clusters were used as sender cells and macrophages, fibroblasts or endothelial cells were set as receiver cells. The condition for receiver cells were transcripts upregulated after myocardial infarction, compared to sham hearts. The top 200 targets and top 30 ligands expressing in at least 10% of each TCR-M cluster were set for the analysis. The obtained results were depicted as Circos plots.

*Statistical analyses*

The results are shown as the mean ± the standard error of the mean (SEM) along with the distribution of all individual values in each group.  Sample sizes for each group is described in figure legends. Graphs and statistical analyses were performed with GraphPad Prism (version 7.0, GraphPad Software, San Diego, CA, USA). Unpaired two-tailed *t*-test was used for comparison between two groups with data following normal distribution. For multiple comparisons between more than two groups, one or two-way analyses of variance (ANOVA) were conducted followed by *post hoc* test. Differences were considered significant for P values below 0.05.

Supplemental Figures 1-5

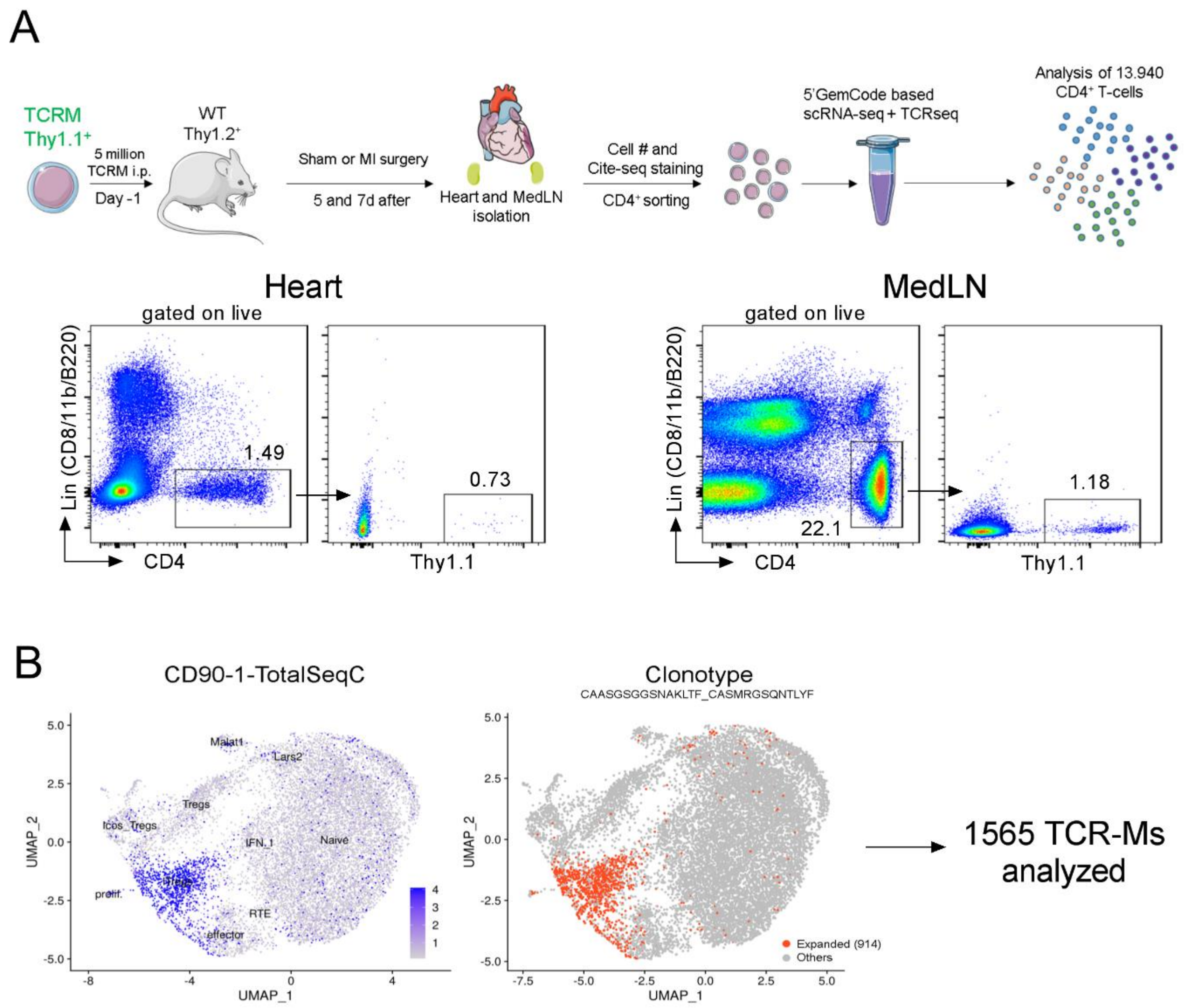


**Supplemental Figure 1**. Single cell RNA sequencing of myosin specific CD4+ T-cells after MI. **(A)** Experiment design: TCR-M cells were transferred one day prior to experimental myocardial infarction to WT Thy1.2 mice. Heart and MedLN from sham operated and infarcted mice at 5 and 7 days post-surgery were simultaneously collected stained with hashtag antibodies (TotalSeqC) and Cite-seq antibodies for CD25 (clone PC61) and CD90.1 (clone OX-7). Heart and MedLNs from all conditions were pooled and Lin-CD4+ cells were sorted in RPMI media plus 20% FCS. Cells were then washed and resuspended with PBS 0.04% BSA. After data cleaning, a total of 13.940 CD4+ T-cells were obtained for analysis. **(B)** scRNAseq data represented in UMAP plots showing TCR-M cells by CD90-1-TotalSeqC signal (left) and by expanded clonotype analysis (914 clones). In total, 1565 TCR-M cells were obtained for analysis.

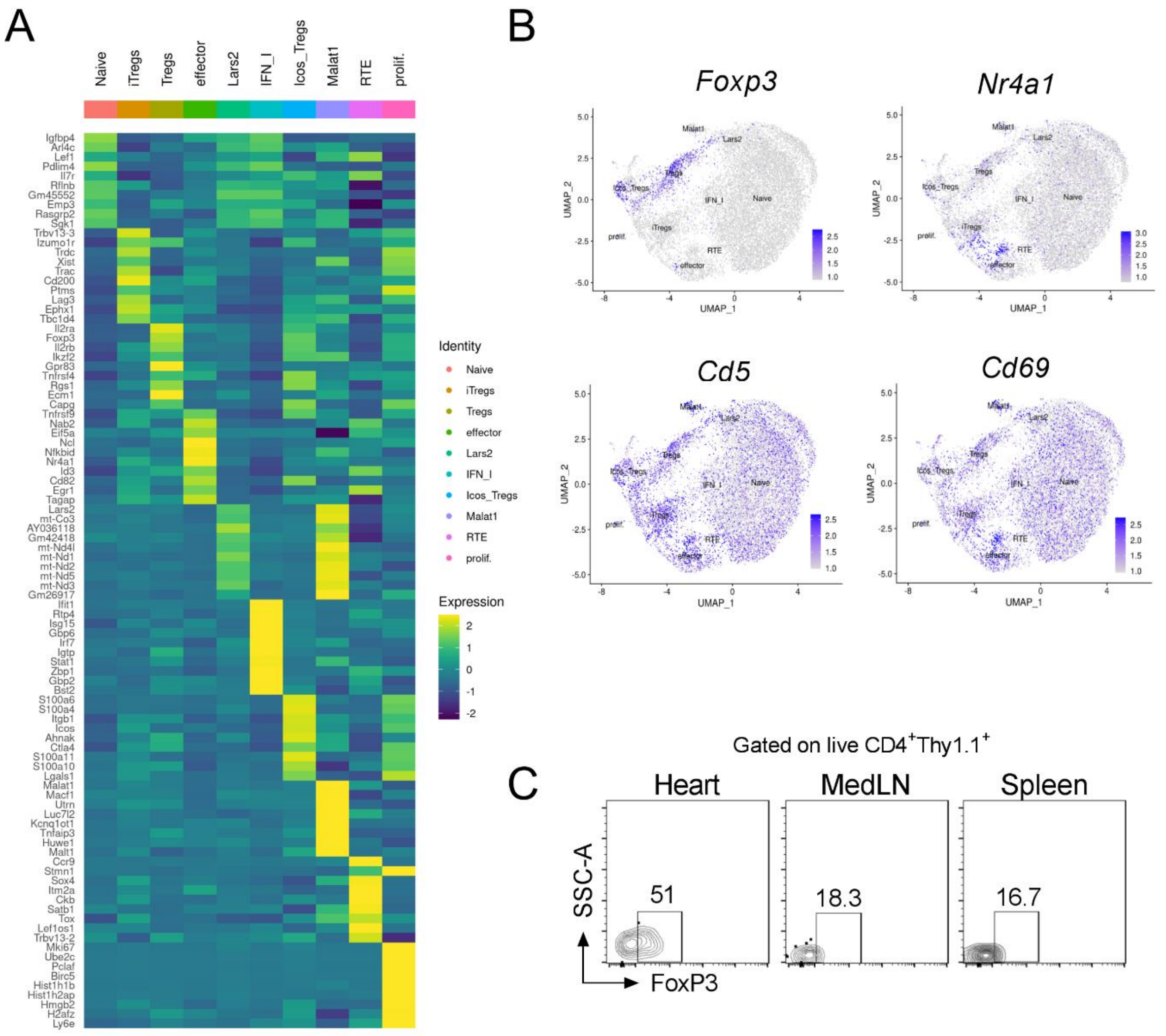


**Supplemental Figure 2**. Expression of cluster defining transcripts in heart and MedLN TCR-M cells and CD4+ endogenous T-cells. **(A)** Heatmap illustrates the top 10 most expressed transcripts in each cell clusters, as defined in Figure 1C. **(B)** *Featureplots* depicting *Foxp3*, *Nr4a1* and pro-activation markers (*Cd5, Cd69*) expression in CD4+ T-cells. (**C**) Representative contour-plots illustrate FOXP3 expression in TCR-M cells isolated from heart, MedLN and spleen at 7d post-MI.

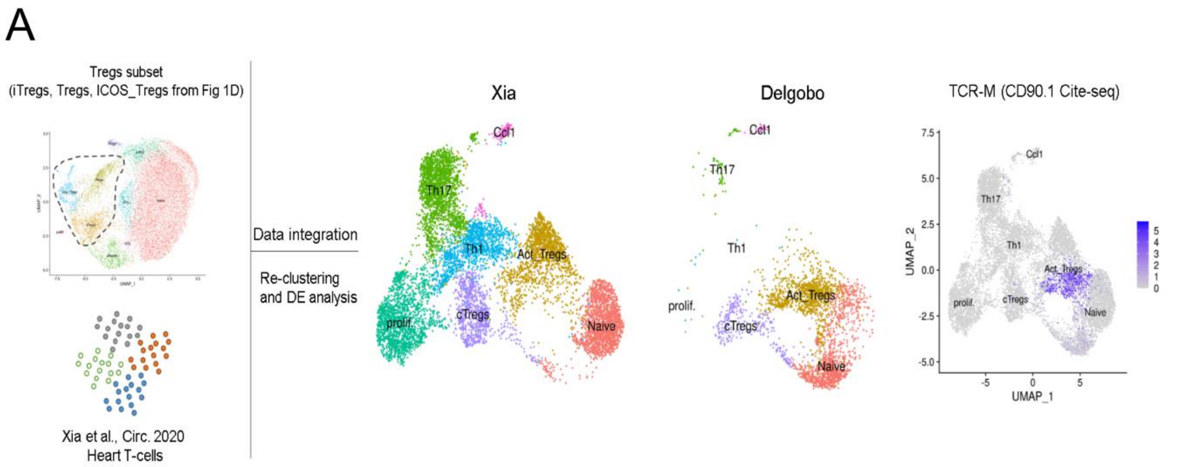


**Supplemental Figure 3**. Distinct TCR-M Treg cluster. **(A)** Data integration with Xia et al study [5]; *iTregs*, *Tregs* and *Icos_Tregs* were subset accordingly (**Figure 1C**) and integrated with Heart T-cell data from infarcted mice at 7d post-surgery. Data was normalized and clusters containing $CD8^+$ T-cells, NK cells, fibroblasts macrophages and B-cells were excluded from combined object. Finally, combined object was restricted to $CD4^+$ T-cells and clusters were classified according to prototypic markers, literature assessment and previously determined cluster signatures from figure 1C (e.g. *Rorc, Il17a, Il17f* for $T_H17$; *Stat1, Tbx21, Itgb1* for $T_H1$). UMAP plots split by study (Xia on the left and Delgobo on the right) reinforce the regulatory profile of TCR-Ms obtained from our study. Right panel shows that TCR-M cells from combined object do not cluster with cells in $T_H17$, $T_H1$ and nor classical Tregs (cTregs).

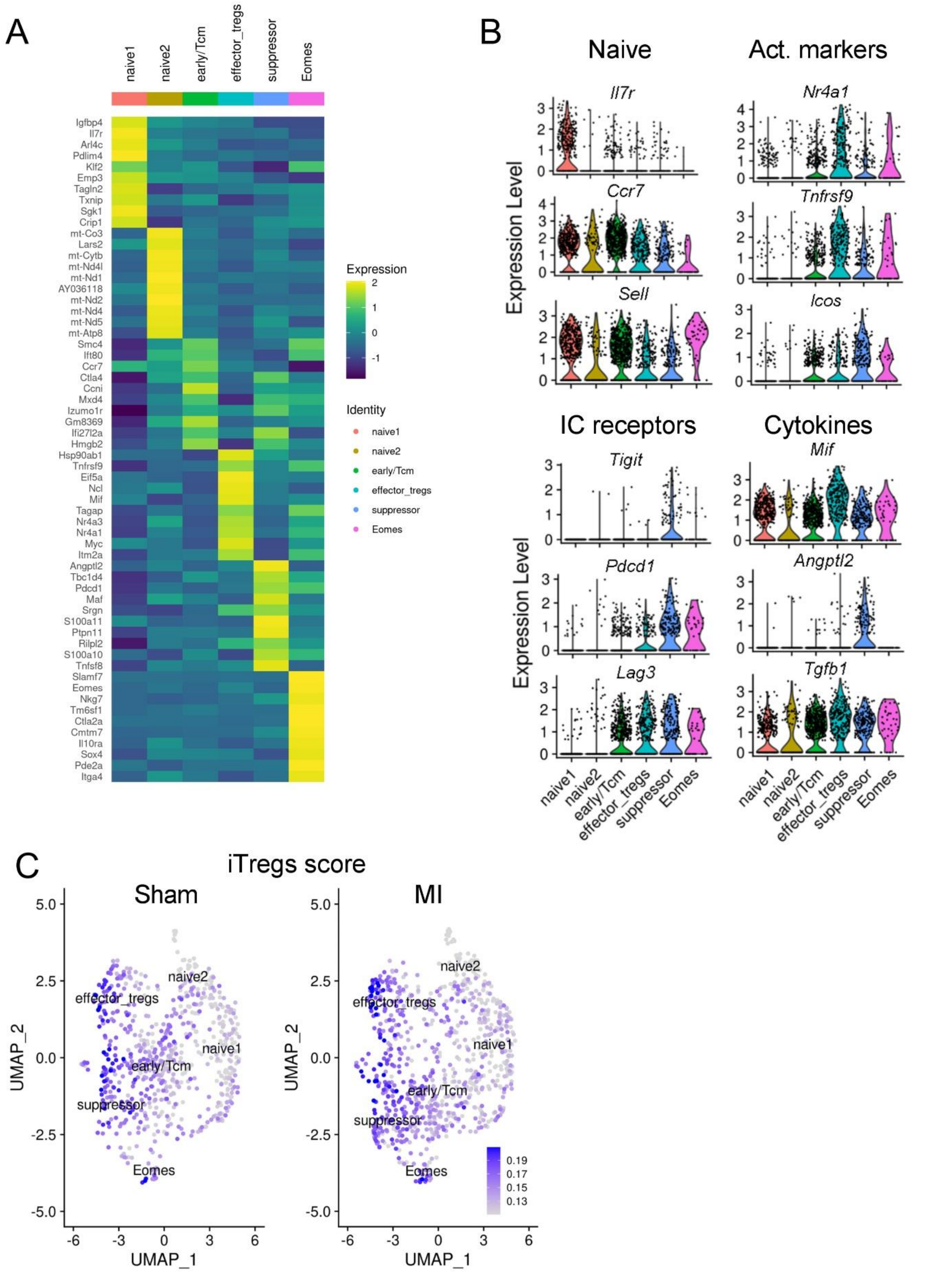


**Supplemental Figure 4**. Transcriptional signature of TCR-M cell clusters. **(A)** Heatmap illustrates the top 10 most expressed transcripts by cell clusters from figure 2A. **(B)** Violin plots showing the expression of genes involved in activation states (Naïve/Act. Markers) and T-cell phenotype/function (IC receptors/Cytokines) at different TCR-M clusters. **(C)** Feature plots from figure 2B illustrates *iTreg* gene module score in TCR-M cells from sham operated or infarcted mice.

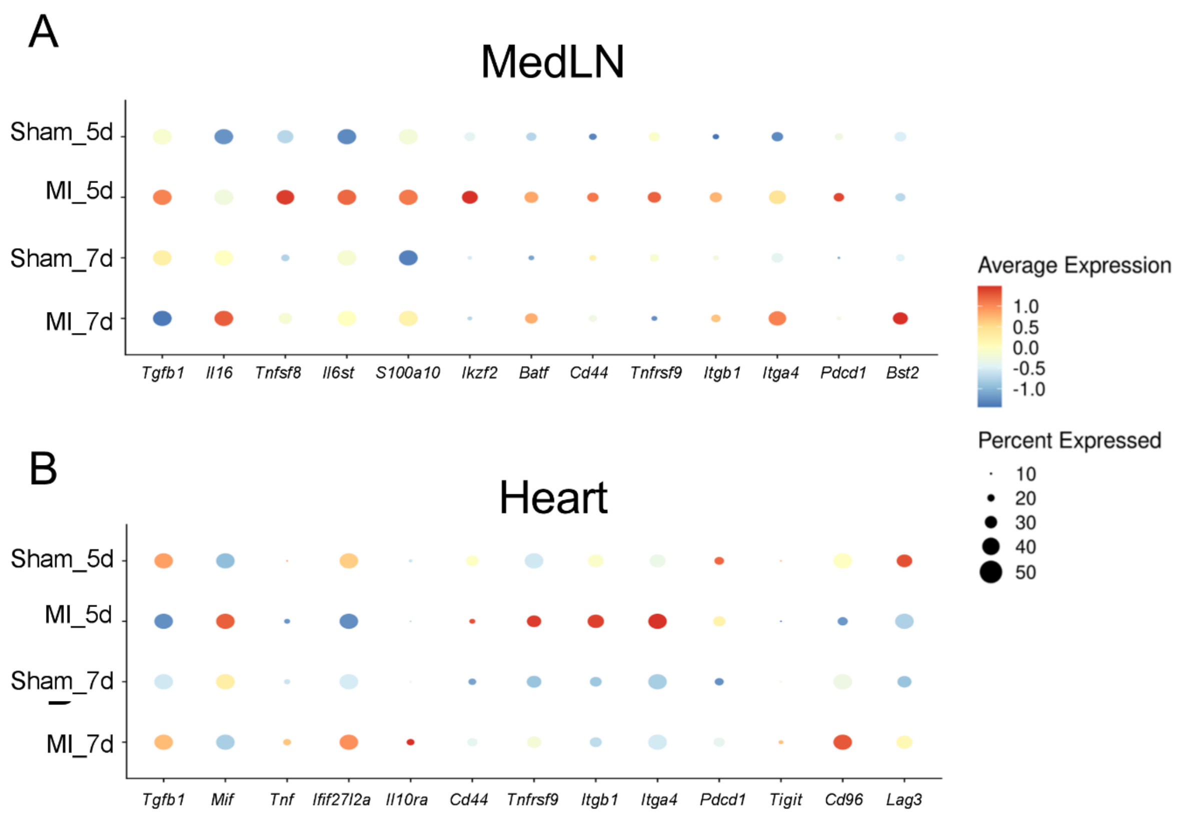


**Supplemental Figure 5**. Expression of cytokines, transcription factors and activation markers in MedLN (**A**) and cardiac (**B**) TCR-M cells at different conditions. (**A**) Average expression value is depicted in heatmap scale and circle size represents percentage expressed.

**Supplementary Table 1 –** Echocardiography measurements in control non-transferred and $T_H$ transferred TCR-M mice at day 5 post-MI surgery.

| | **Control** | **$T_H$1** | **$T_H$17** | **Treg** |
|---|---|---|---|---|
| Survival (%) | 95.2 | 78.6 | 84.2 | 92.3 |
| **Heart Rate (bpm)** | 435.9 ± 15.94 | 455.2 ± 29.61 | 472.24 ± 16.25 | 436.7 ± 13.6 |
| **Infarct size (%)** | 36.96 ± 2.13 | 40.92 ± 0.89 | 36.55 ± 2.31 | 30.57 ± 3.125 |
| **Ejection fraction (%)** | 35.99 ± 2.65 | 45.88 ± 2.69 | **45.66 ± 3.15*** | 41.81 ± 2.11 |
| **EDA ($mm^2$)** | 15.45 ± 1.05 | 12.82 ± 0.87 | 12.69 ± 1.31 | **11.11 ± 1.06*** |
| **ESA ($mm^2$)** | 12.63 ± 1.19 | 9.04 ±1.088 | **8.20 ± 1.4*** | **7.37 ± 0.75*** |

N: 6-11 per group. Statistical analysis: One-way ANOVA followed by Dunnett's post-hoc test, *$p<0.05$ against control group. Survival analysis: Log-rank test, p:0.44. EDA: End diastolic area, ESA: End systolic area.

References


1. Nindl V, Maier R, Ratering D, De Giuli R, Zust R, Thiel V, Scandella E, Di Padova F, Kopf M, Rudin M, Rulicke T, Ludewig B. Cooperation of th1 and th17 cells determines transition from autoimmune myocarditis to dilated cardiomyopathy. *Eur J Immunol*. 2012;42:2311-2321
2. Hofmann U, Beyersdorf N, Weirather J, Podolskaya A, Bauersachs J, Ertl G, Kerkau T, Frantz S. Activation of cd4+ t lymphocytes improves wound healing and survival after experimental myocardial infarction in mice. *Circulation*. 2012;125:1652-1663
3. Lindsey ML, Bolli R, Canty JM, Jr., Du XJ, Frangogiannis NG, Frantz S, Gourdie RG, Holmes JW, Jones SP, Kloner RA, Lefer DJ, Liao R, Murphy E, Ping P, Przyklenk K, Recchia FA, Schwartz Longacre L, Ripplinger CM, Van Eyk JE, Heusch G. Guidelines for experimental models of myocardial ischemia and infarction. *Am J Physiol Heart Circ Physiol*. 2018;314:H812-H838
4. Farbehi N, Patrick R, Dorison A, Xaymardan M, Janbandhu V, Wystub-Lis K, Ho JW, Nordon RE, Harvey RP. Single-cell expression profiling reveals dynamic flux of cardiac stromal, vascular and immune cells in health and injury. *Elife*. 2019;8
5. Xia N, Lu Y, Gu M, Li N, Liu M, Jiao J, Zhu Z, Li J, Li D, Tang T, Lv B, Nie S, Zhang M, Liao M, Liao Y, Yang X, Cheng X. A unique population of regulatory t cells in heart potentiates cardiac protection from myocardial infarction. *Circulation*. 2020;142:1956-1973
